# E2F1 Mediates SOX17 Deficiency-Induced Pulmonary Hypertension

**DOI:** 10.1101/2023.02.15.528740

**Authors:** Dan Yi, Bin Liu, Hongxu Ding, Shuai Li, Rebecca Li, Jiakai Pan, Karina Ramirez, Xiaomei Xia, Mrinalini Kala, Indrapal Singh, Qinmao Ye, Won Hee Lee, Richard E. Frye, Ting Wang, Yutong Zhao, Kenneth S. Knox, Christopher C. Glembotski, Michael B. Fallon, Zhiyu Dai

## Abstract

**Rationale:** Rare genetic variants and genetic variation at loci in an enhancer in SRY-Box Transcription Factor 17 (SOX17) are identified in patients with idiopathic pulmonary arterial hypertension (PAH) and PAH with congenital heart disease. However, the exact role of genetic variants or mutation in SOX17 in PAH pathogenesis has not been reported.

**Objectives:** To investigate the role of SOX17 deficiency in pulmonary hypertension (PH) development.

**Methods:** Human lung tissue and endothelial cells (ECs) from IPAH patients were used to determine the expression of SOX17. Tie2Cre-mediated and EC-specific deletion of Sox17 mice were assessed for PH development. Single-cell RNA sequencing analysis, human lung ECs, and smooth muscle cell culture were performed to determine the role and mechanisms of SOX17 deficiency. A pharmacological approach was used in Sox17 deficiency mice for therapeutic implication.

**Measurement and Main Results:** SOX17 expression was downregulated in the lungs and pulmonary ECs of IPAH patients. Mice with Tie2Cre mediated Sox17 knockdown and EC-specific Sox17 deletion developed spontaneously mild PH. Loss of endothelial Sox17 in EC exacerbated hypoxia-induced PH in mice. Loss of SOX17 in lung ECs induced endothelial dysfunctions including upregulation of cell cycle programming, proliferative and anti-apoptotic phenotypes, augmentation of paracrine effect on pulmonary arterial smooth muscle cells, impaired cellular junction, and BMP signaling. E2F Transcription Factor 1 (E2F1) signaling was shown to mediate the SOX17 deficiency-induced EC dysfunction and PH development.

**Conclusions:** Our study demonstrated that endothelial SOX17 deficiency induces PH through E2F1 and targeting E2F1 signaling represents a promising approach in PAH patients.

## Introduction

A population-based study involving 3,381 people suggests that the prevalence of echocardiographic signs of possible pulmonary hypertension (PH) is 2.6% of the general population(1, 2). Heritable and idiopathic PAH (IPAH), also previously known as primary PH, are a form of PH. They are clinically identical progressive disorders charactered by elevation of pulmonary arterial pressure with pathologic remodeling in pulmonary arteries(3). PH types with different etiologies share histopathologic features including eccentric and obliterative intima thickening and complex plexiform lesions. *BMPR2*, a gene encoding bone morphogenetic protein type 2 receptor (BMPR2), is mutated in 80% of familial PAH and approximately 20% of sporadic cases. Other mutations or pathogenic genes have been identified, including other TGF-β/BMP signaling members *ACVRL1, ENG, SMAD1/4/9, and CAV1, KCNK3*, and *TBX4*(4). Recent studies also identified a few rare sequence variations in the genes *GDF2, ATP13A3, AQP1*, and *SOX17*(5). However, the exact mechanisms by which these gene mutations or variants increase the susceptibility to PH remain elusive.

A transcription factor SOX17, a member of the Sry-related high mobility group domain family F (Sox F) transcription factors, is a critical regulator in the developmental stage of endothelial/hematopoietic lineages and maintenance of arterial identities(6–8). In the developmental lung, SOX17 is selectively expressed in the pulmonary arteries and veins. Interestingly, SOX17 is only detected in the vasculature of the right ventricle in the developmental heart(9). Deletion of Sox17 (Sox17^Δ/Δ^) at embryonic stage causes pulmonary vascular malformations, biventricular enlargement and postnatal lethality(10), suggesting that endothelial SOX17 is critical to cardiopulmonary development. In the adult lung, Sox17 is required for endothelial regeneration following sepsis-induced vascular injury in mice(11). Endothelial SOX17 also promotes tumor angiogenesis(12). Rare genetic variants in *SOX17* are identified in patients with IPAH and PAH with congenital heart disease (CHD)(13, 14). Recent studies also identified genetic variation at loci in an enhancer near SOX17 is associated with PAH(15). A recent study showed that Sox17 deficiency promoted PH in mice via HGF/c-Met signaling(16). Nevertheless, the exact role of genetic variants or mutation in SOX17 in the contribution of PH remain unclear.

In our present studies, we showed that SOX17 is downregulated in pulmonary arterial endothelial cells (PAECs) isolated from IPAH patients compared to healthy donors. Using EC-specific deletion mouse model, we demonstrated, for the first time, that deficiency of Sox17 in ECs in mice induced spontaneously vascular remodeling and mild PH, and augmented hypoxia-induced PH. Loss of SOX17 in human PVECs (HPVECs) stimulated EC hyperproliferation and apoptosis resistance, which is likely due to the activation of transcriptional factor E2F1 and its downstream programming. Targeting E2F1 signaling represents an effective approach for inhibiting SOX17 deficiency-induced vascular remodeling in PAH patients.

## Materials and Methods

### Human samples

The use of archived human lung tissues and cells were granted by the University of Arizona (UA) Institutional Review Board. Human IPAH patients and failed donors (FD)’ PVECs were obtained from the Pulmonary Hypertension Breakthrough Initiative (PHBI).

### Mice

CKO Sox17 mice were generated by breeding *Sox17*^f/f^ mice with Tie2Cre mice(17). ecKO *Sox17* mice were generated by breeding *Sox17*^f/f^ mice with *EndoSCL-CreERT2* mice(18). Both male and female mice were included for experiments. For HLM treatment, ecKO *Sox17* mice were treated with tamoxifen, followed by treatment with HLM006474 (HLM, 12.5 mg/kg) 3 times a week for 6 weeks. The protocol for animal care and studies was approved by the Institutional Animal Care and Use Committee of UA.

### Data availability

RNA-seq and scRNA-seq data have been deposited in the GEO database under accession number GSE192649. Scripts used for single-cell RNA sequencing analysis and analyzed data in R objects are available in Figshare (https://figshare.com/s/37782988b8cac7cedcf9).

### Statistical Analysis

Statistical determination was performed on Prism 9 (Graphpad Software Inc.). Two-group comparisons were compared by the unpaired 2-tailed Student t test for equal variance or the Welch t test for unequal variance. Multiple comparisons were performed by One Way ANOVA with a Tukey post hoc analysis that calculates corrected P values. P less than 0.05 indicated a statistically significant difference. All bar graphs represent mean±SD.

## Results

### SOX17 is downregulated in PVECs from PAH patients

SOX17 mutations and enhancer variants were found in patients with PAH. However, the expression pattern and levels of SOX17 in human PAH patients remain elusive. Leveraging the public single-cell RNA-sequencing dataset from healthy human lungs, we first analyzed the mRNA expression of *SOX17*. Our data demonstrated that *SOX17* is highly expressed in the endothelial cells (ECs) and rarely expressed in other cell types in the adult lung (**Figure 1A**). To determine whether SOX17 is deficient in PVECs of PAH patients, we characterized the SOX17 expression in isolated PVECs from IPAH patients and failed donors (FD). We found that the SOX17 mRNA levels (**Figure 1B**) as well as the SOX17 protein levels (**Figure 1C**) were significantly downregulated in sub-confluent PVECs isolated from IPAH patients compared to that from FD subjects, suggesting that SOX17 deficiency is present in PAH patients. Our data is consistent the microarray analysis of lung samples from IPAH patients and healthy donors, which showed that SOX17 mRNA level is decreased in IPAH patients(19) (**Supplemental Figure 1A**). To determine the localization of SOX17, we performed immunofluorescent staining against SOX17 on human IPAH and FD lungs and our data showed that SOX17 is mainly located in the lung ECs. As shown in **Figure 1D, 1E and Supplemental Figure 1B**, SOX17 is markedly downregulated in the ECs of less remodeled vessels and diminished in the occlusive vessels of IPAH patients. We also determine the levels of SOX17 in the lung of monocrotaline (MCT) induced PH rats, we found that there was a significant reduction of SOX17 in MCT-treated rats (**Figure 1F**).

**Figure 1.**
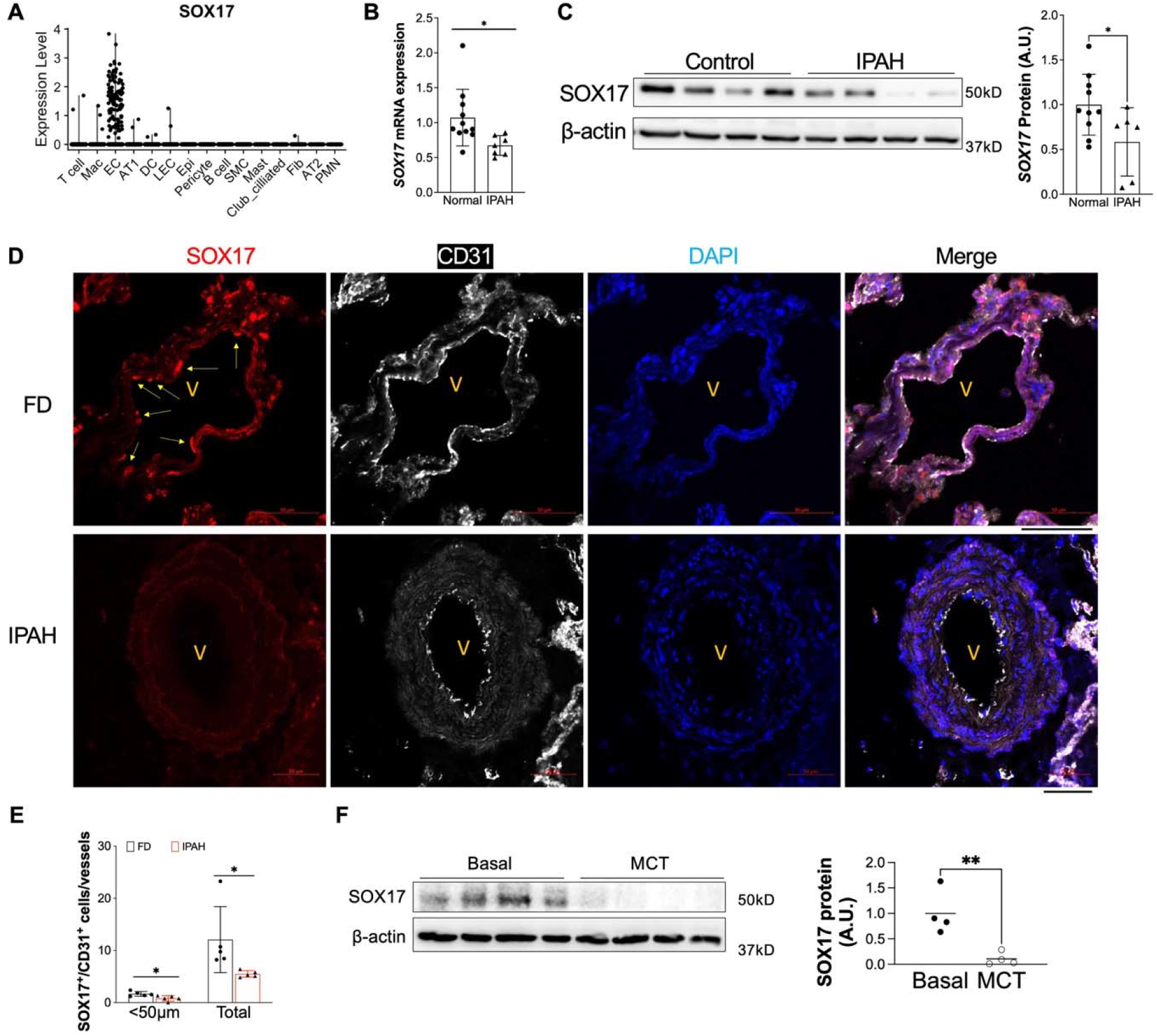
Downregulation of endothelial SOX17 in the patients with PAH. (A) A violin plot showing SOX17 is restricted in the ECs of human lungs via scRNA-seq. Mac=macrophage; DC= dendritic cell; LEC=lymphatic EC; Epi=epithelium; SMC= smooth muscle cell; Fib=fibroblast; AT1 or AT2 = alveolar type 1 or 2 epithelium; PMN=neutrophils. (B) qRT-PCR analysis showed that SOX17 mRNA levels were downregulated in the sub-confluent PVECs isolated from IPAH patients. Each data point represents cells from one human subject including both male and female. (C) Western blotting demonstrated reduction of SOX17 protein expression in the IPAH PVECs. Each data point represents cells from one human subject including both male and female. (D, E) Immunostaining against SOX17 showing diminished SOX17 expression in the ECs of remodeling lesions from IPAH patients. Arrows indicate SOX17 positive ECs in non-PAH failed donors (FD). SOX17^+^/CD31^+^ cell number was quantified and normalized by vessels number. Each dot represents one subject. (F) SOX17 is decreased in the lungs of established PH rats at 4 weeks post MCT (33mg/kg subcutaneously) treatment. Student t test (B, C, E, F). *, P< 0.05; **, P< 0.01. A.U. = arbitrary units; Scale bar, 50*μ*m.

### Loss of SOX17 in embryonic stage induces spontaneously mild PH and cardiac hypertrophy

To determine whether SOX17 deficiency is involved in the pathogenesis of PH in mice, we utilized EC specific Cre lines (Tie2Cre and EndoSCL-CreER^T2^) to delete Sox17 in the ECs. Constitute deletion of Sox17 mice (Sox17^f/f^;Tie2Cre) display vascular defect and embryonic lethal(10), thus we generated Sox17^f/+^;Tie2Cre (KO^*EC+*^, cKO) mice. We then characterized the right ventricular (RV) hemodynamic and cardiac dissection of WT (Sox17^f/f^) and cKO mice. cKO mice at the basal developed mild PH by upregulation of right ventricle systolic pressure (RVSP), which is the indicator of pulmonary arterial pressure, when compared with WT mice in the similar age (**Figure 2A**). We also observe a significant increase in the weight ratio of the right ventricular free wall to left ventricle plus septum (RV/LV+S) and left ventricle weight vs body weight (LV/BW), indicative of right ventricular and left ventricular hypertrophy, in cKO mice. (**Figure 2B and 2C**), which is consistent with previous finding that embryonic deletion of Sox17 lead to enlargement of biventricles (10). To further determine whether Tie2Cre promoter mediated Sox17 knockdown regulates pulmonary vascular remodeling in mice, we performed Russell-Movat pentachrome staining and immunostaining of α-smooth muscle actin (SMA) and found that cKO mice exhibited increased of the thickness of pulmonary arterial wall, and muscularization of distal pulmonary arterioles (**Figure 2D and 2F**).

**Figure 2.**
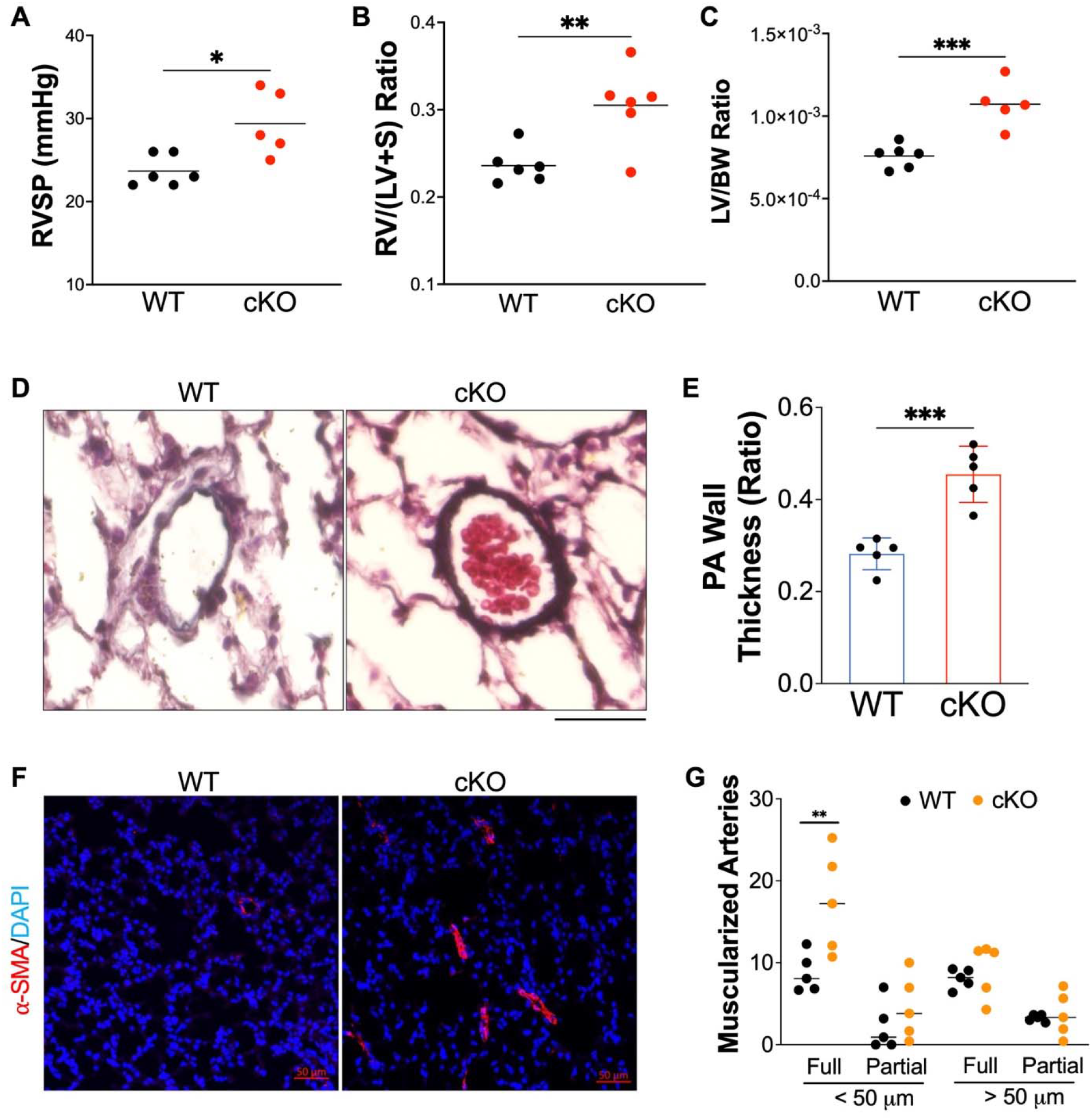
Tie2Cre mediated Sox17 deficiency induced PH and cardiac hypertrophy. (A) Hemodynamic measurement showing that cKO Sox17 mice had increased right ventricular systolic pressure (RVSP) compared with *Sox17^f/f^* (WT) mice. (B and C) Cardiac dissection showed the upregulation of right heart and left heart hypertrophy in cKO mice compared with WT mice.. (D) Representative micrographs of Russell-Movat pentachrome staining showing increased medial thickness in Sox17 cKO mice compared with WT mice. (E) Quantification of pulmonary artery wall thickness. Wall thickness was calculated by the distance between internal wall and external wall divided by the distance between external wall and the center of lumen. (F and G) Muscularization of distal pulmonary vessels was markedly enhanced in Sox17 cKO mice compared with WT mice. Lung sections were immunostained with anti–*α*-SMA (green). Red arrow indicates a-SMA^+^ distal pulmonary vessels. α-SMA^+^ vessels were quantified in 20 field at 10X magnification per mouse (D) Student t test (A, B, C, E, G). *, P< 0.05; **, P< 0.01, ***, P< 0.001. Scale bar, 50*μ*m.

### Loss of endothelial SOX17 in adult stage leads to spontaneously mild PH

Because Tie2Cre also induces gene deletion in hematopoietic stem cells besides ECs(20), we then generated inducible deletion of EC Sox17 mice (*Sox17*^f/f^;EndoSCL-CreERT2(18, 21), ecKO *Sox17*) by breeding *Sox17* floxed mice with EndoSCL-CreERT2(18, 21) (**Supplemental Figures 2A**). Both *Sox17*^f/f^ (WT) and ecKO *Sox17* mice at the age of 7~8 weeks were treated with tamoxifen for 3 doses [100mg/kg, intraperitoneal injection (i.p.) daily] to induce SOX17 deletion only in ECs. Around 2 months post tamoxifen treatment, Immunostaining against SOX17 demonstrated that PVECs from ecKO *Sox17* mice have significant decrease of SOX17 expression, suggest that SOX17 was selectively deleted in PVECs (**Supplemental Figures 2B**). We then characterized the RV hemodynamic and cardiac dissection of WT and ecKO *Sox17* mice. Our data showed that ecKO *Sox17* mice showed a significant increase of RVSP when compared with WT mice (**Figure 3A)**. However, we did not observe a significant change in RV/LV+S ratio and LV/BW ratio between WT and ecKO *Sox17* mice (**Figure 3B and 3C**). We also performed echocardiography measurement on these animals. We did not observe any significant alteration of cardiac size and function including heart rate, cardiac output, left ventricular fractional shorting and RV fraction area change in the ecKO Sox17 mice (**Supplemental Figure 2C-2F**). The difference cardiac phenotype between Sox17 cKO and ecKO mice might be due to the effect of constitute Sox17 deletion in the embryonic stage. To further determine whether endothelial Sox17 deficiency regulates pulmonary vascular remodeling in mice, we then performed Russell-Movat pentachrome staining (**Figure 3D and 2E**) and immunostaining of α-smooth muscle actin (SMA) (**Figure 2F and 2G**). Examination of lung pathology showed that ecKO *Sox17* mice exhibited a marked increase of pulmonary wall thickness and distal pulmonary arterial muscularization assessed by α-SMA staining (**Figure 3, D-G**), demonstrating loss of endothelial SOX17 aggravates pulmonary vascular remodeling in mice. As PAH is associated with upregulation of accumulation of perivascular inflammatory, we found that ecKO *Sox17* mice exhibited increased CD45^+^cells accumulation in the vascular bed compared to WT mice (**Figure 3H and 3I**). Taken together, our data demonstrated that Sox17 deficiency induces PH in mice.

**Figure 3.**
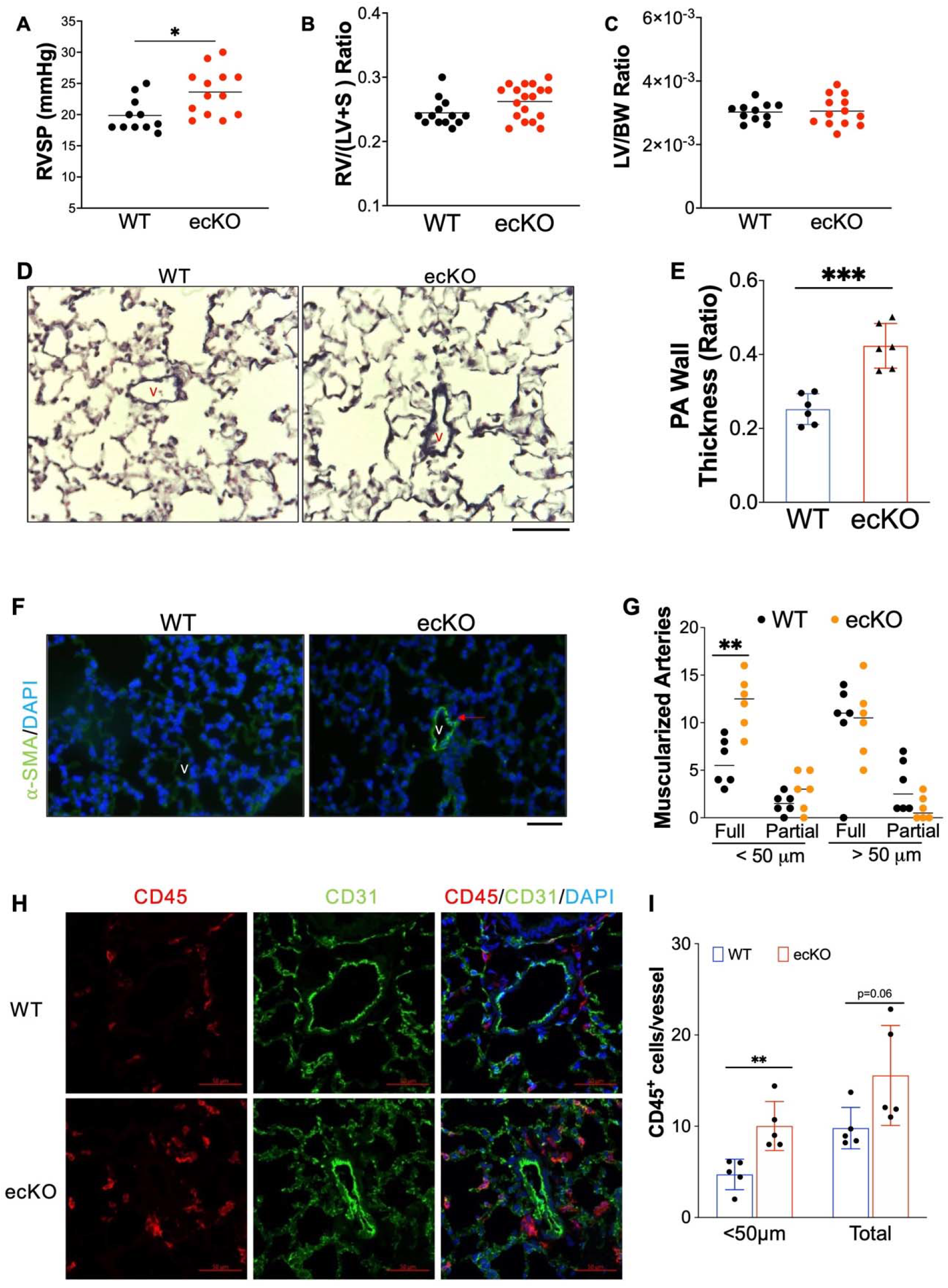
Endothelial SOX17 deficiency induced spontaneous mild PH. (A) ecKO Sox17 mice exhibited increase of RVSP. (B and C) No change of RV and LV hypertrophy in ecKO Sox17 mice compared with WT mice. (D) Representative micrographs of Russell-Movat pentachrome staining showing increased medial thickness in ecKO Sox17 mice compared with WT mice. (E) Quantification of pulmonary artery wall thickness. Wall thickness was calculated by the distance between internal wall and external wall divided by the distance between external wall and the center of lumen. (F and G) Muscularization of distal pulmonary vessels was markedly enhanced in ecKO Sox17 mice compared with WT mice. Lung sections were immunostained with anti–α-SMA (green). Red arrow indicates a-SMA^+^ distal pulmonary vessels. α-SMA^+^ vessels were quantified in 20 field at 10X magnification per mouse. (H and I) Immunostaining against CD45 (Red) demonstrated that there was upregulated accumulation of inflammatory cells in the perivascular bed of ecKO Sox17 mice. Student t test (A, B, C, E, G, I). *, P< 0.05; **, P< 0.01, ***, P< 0.001. Scale bar, 50*μ*m.

### Loss of SOX17 in ECs exaggerated hypoxia-induced PH

Previous studies demonstrated that Sox17 is a HIF-1α target gene in the lung ECs(11). We did find that Sox17 is upregulated in the lung of chronic hypoxia incubated mice (**Supplemental Figure 3**). To further confirm if SOX17 deficiency in EC augments PH and RV remodeling in mice, we challenged both WT and ecKO *Sox17* mice with hypoxia (10% O_2_) to assess the role of endothelial ecKO *Sox17* in the hypoxia-induced PH in mice. Both WT and ecKO *Sox17* mice at the age of 7~8 weeks were treated with tamoxifen for 3 doses (20mg/kg, i.p. injection daily) to induce SOX17 deletion. 3 weeks post tamoxifen treatment, mice were incubated with hypoxia (10% O_2_) for 3 weeks or normoxia alone. Our data showed that ecKO *Sox17* mice exposed to hypoxia exhibited a significantly elevated of RVSP when compared with WT mice (**Figure 4A**). ecKO *Sox17* mice also showed a significantly increased weight ratio of RV/(LV+S), indicative of RV hypertrophy compared with WT mice (**Figure 4B**). We then examined the pulmonary pathology and found the narrower pulmonary vessel lumen and thicker wall in the big vessels of ecKO *Sox17* mice (**Figures 4C** and **4D**). In addition, we also observed occasional occlusion in the small vessels of ecKO *Sox17* mice but not in WT mice (**Figure 4C**). Moreover, there is an increased muscularization of distal pulmonary arteries in the ecKO *Sox17* mice compared with WT mice (**Figures 4E-4F**). These studies showed that genetic deletion of endothelial SOX17 augmented hypoxia-induced pulmonary vascular remodeling and vasoconstriction in mice.

**Figure 4.**
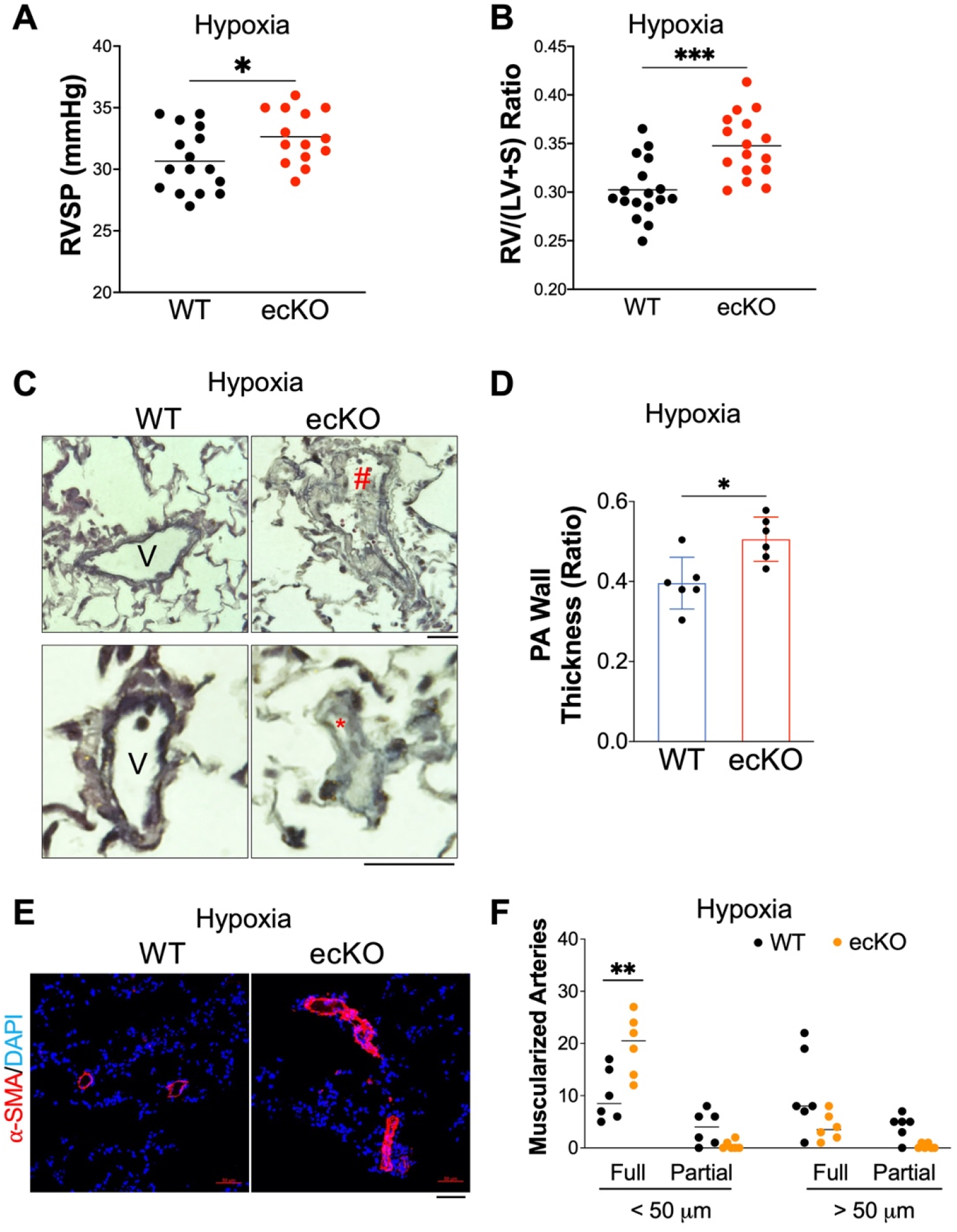
Augmentation of PH by SOX17 deficiency in ECs under hypoxia. (A) Hemodynamic measurement demonstrated that ecKO Sox17 mice exhibited increased of RVSP compared to WT mice under hypoxia condition. (B) RV dissection showing upregulation of RV hypertrophy in ecKO Sox17 mice compared to WT mice in response to hypoxia. (C and D) Quantification of Russell-Movat pentachrome staining showing thicker pulmonary artery walls and representative micrographs in ecKO Sox17 mice compared with WT mice in hypoxia condition. V=vessel, # indicates narrower vessel, * indicates occlusive vessel. Wall thickness was calculated by the distance between internal wall and external wall divided by the distance between external wall and the center of lumen. (E and F) Quantification of anti–*α*-SMA staining showing upregulation of muscularization of distal pulmonary artery wall and representative micrographs in ecKO Sox17 mice compared with WT mice in hypoxia condition. α-SMA^+^ vessels were quantified in 20 field at 10X magnification per mouse. Student t test (A, B, D and F). *, P< 0.05; **, P< 0.01, ***, P< 0.001. Scale bar, 50*μ*m.

### SOX17 deficiency induces endothelial cell proliferation

To validate the impact of Sox17 deletion in vivo, we applied single-cell RNA sequencing (scRNA-seq) analysis on cKO mice and WT mice (**Supplemental Figure 4A**). scRNA-seq data revealed an increase of EC proportion in cKO mice compared with WT mice (**Supplemental Figure 4B**). Transcriptomic analysis demonstrated that the lung ECs from cKO mice exhibited increased expression of genes related to cell proliferation, including *Cdk1, E2f1, Top2a*, etc (**Figure 5A**). To understand the direct impact of SOX17 deficiency in pulmonary EC in vitro, we also performed whole transcriptome RNA-sequencing in HPVECs with SOX17 knockdown. siRNA against SOX17 efficiently reduced SOX17 mRNA level and proteins expression (**Figures 5B and 5C**). RNA-seq analysis and pathway enrichment analysis showed that there was an alteration of many genes (i.e., *CENPP, BRCA2, CDKN2C, CCNB2*) and pathway (i.e., cell cycle) related to cell proliferation (**Figures 5D** and **5E**). QRT-PCR analysis confirmed that SOX17 knockdown significantly induced expression of genes related to cell proliferation including *PLK1, CCNA2, CCNB1, CCNB2, CDKL1*, and *CKDN2C* (**Figure 5F**). Western Blotting confirmed upregulation of PLK1 protein expression by SOX17 knockdown (**Figure 5G**). As SOX17 deficiency in EC induces cell cycle program, we hypothesize that SOX17 deficiency might lead to endothelial hyperproliferation during the development of PH. We employed siRNA to knockdown SOX17 in cultured HPVECs and evaluated cell proliferation. Cell proliferation, assessed by 5-bromo-2’-deoxyuridine (BrdU) incorporation assay, in SOX17-deficient cells was markedly augmented compared to control siRNA-transfected HPVECs (**Figure 5H)**. We also evaluated in vivo proliferation via injecting BrdU into WT and ecKO mice. We found that BrdU incorporation in CD31^+^ cells were markedly increased in ecKO mice (**Figure 5I**). The expression levels of a cell proliferation marker, proliferating cell nuclear antigen (PCNA), and polo-like kinase 1 (PLK1) were upregulated in the lung from ecKO *Sox17* mice compared to WT mice (**Figure 5J**). These data suggest that SOX17 deficiency induces EC proliferation in vitro and in vivo.

**Figure 5.**
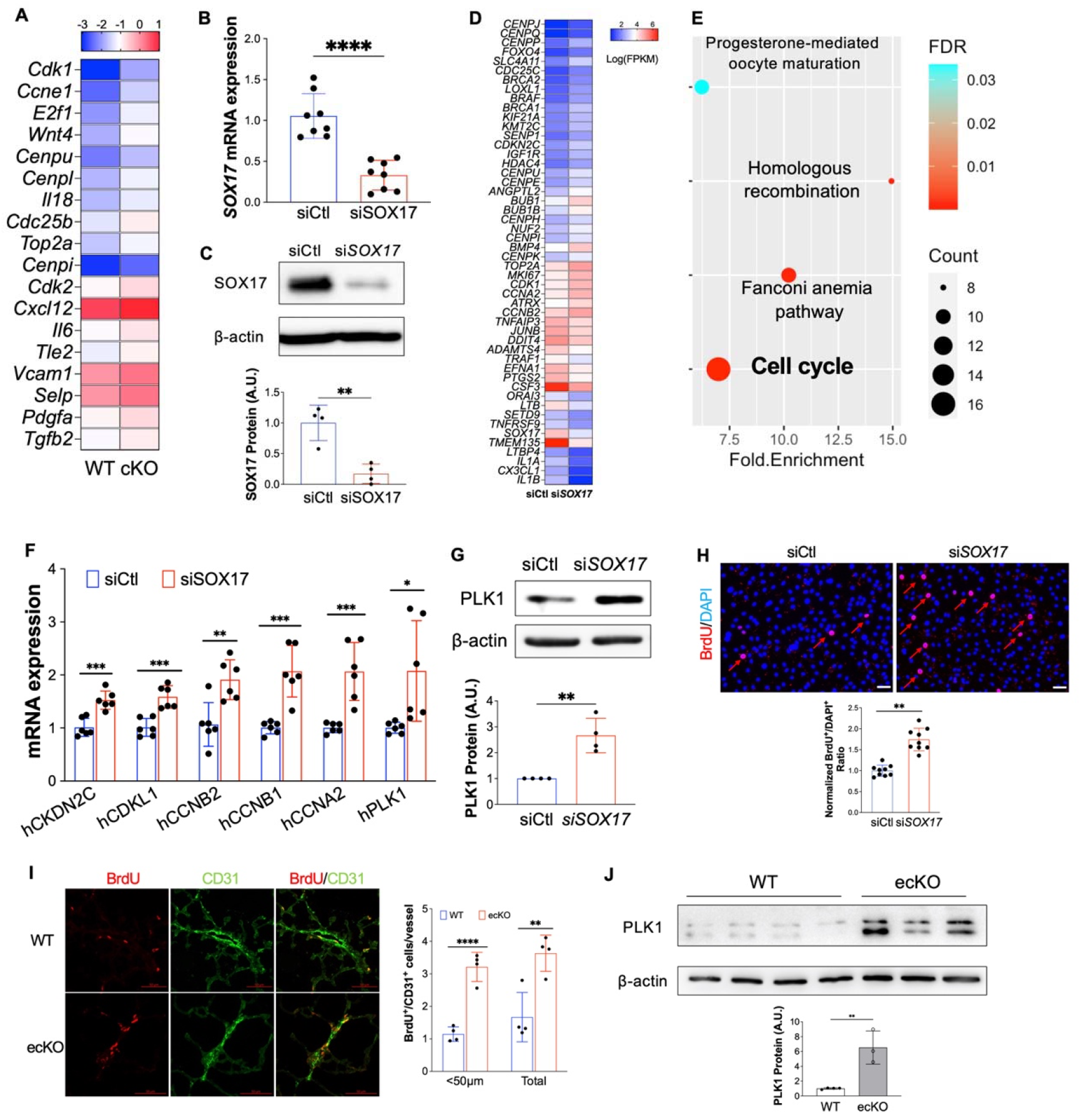
Loss of SOX17 induced EC proliferation. (A) scRNA transcriptomics showed that Sox17 deficiency ECs expressed higher levels of proliferation genes compared to WT ECs. scRNA-seq analysis was performed on the whole lung of WT and cKO mice. Lung ECs transcriptomics were analyzed. (B) qRT-PCR analysis showing efficient knockdown of SOX17 via siRNA against SOX17 in HPVECs. (C) siRNA against SOX17 markedly reduced SOX17 protein expression. (D) A representative heatmap of RNA-sequencing analysis of SOX17 knockdown in HPVECs. HPVECs were transfected with control siRNA (siCtl) or SOX17 siRNA for 48 hours. Equal amount of RNA from three replicates per group were pooled for RNA-seq. (E) KEGG pathway enrichment analysis of upregulated genes in SOX17 deficient lung ECs demonstrating that cell cycle pathway is the top upregulated signaling induced by loss of SOX17. (F) qRT-PCR analysis confirmed the upregulation of cell proliferation related genes including *CKDN2C, CDKL1, CCNB2, CCNB1, CCNA2, and PLK1*. (G) Western Blotting analysis demonstrated induction of PLK1 protein expression by SOX17 deficiency. (H) BrdU incorporation assay demonstrated increased of EC proliferation in SOX17 deficient HPVECs. At 48 hours post-transfection, HPVECs were starved in serum/growth factors free medium for 12 hours. BrdU was added in the medium at 4 hours prior to cells harvest. BrdU was stained with anti-BrdU antibodies. Red indicated BrdU positive cells. Nucleus were co-stained with DAPI. (I) In vivo BrdU incorporation assay showed upregulation of lung ECs proliferation in ecKO Sox17 mice during hypoxia condition. WT and ecKO Sox17 mice were incubated in hypoxia (10% O_2_) for 10 days. BrdU (25 mg/kg) was injected i.p. between day 7 to day 9. Lung sections were stained with anti-BrdU and anti-CD31. BrdU^+^/CD31^+^ cells were quantified. (J) Augmentation of cell proliferation marker PLK1 expression in the lung of ecKO Sox17 (ecKO) mice compared to WT mice. β-actin level was used as an internal control. Student t test (B, C, F, G, H, J). *, P< 0.05; **, P< 0.01. ***, P< 0.001. Scale bar, 50*μ*m.

### Endothelial SOX17 deficiency induces PASMCs proliferation

The muscularization of distal pulmonary arterials and neointima formation seen in ecKO mice are likely due to increased proliferation of pulmonary arterials smooth muscle cell (PASMCs). PAECs from PAH patients produce pro-proliferative signaling through secreting many growth factors such as PDGF-B, ET-1, CXCL12, and MIF, and promote perivascular cells such as PASMCs proliferation(22). We then seeded SOX17 deficient HPVECs on the top chamber and co-cultured with PASMCs, and found that SOX17 knockdown promoted PASMCs proliferation (**Figure 6A-6C**). In vivo BrdU assay also showed the increased of BrdU^+^/α-SMA^+^ cells (indicating PASMCs) proliferation in the Sox17 ecKO mice compared to WT mice (**Figure 6D-6E**). These data suggest that SOX17 deficiency induces paracrine effect and enhances PASMCs proliferation. To identify the potential factors derived from SOX17 deficiency ECs, we leveraged the scRNA-seq dataset and predicted the potential ligand and receptor pairs between ECs and SMCs using CellChat(23). CellChat prediction showed that there were increased ligand-receptor pairs such as Pdgfb-Pdgfra, Edn1-Ednra from ECs to PASMCs (**Figure 6F**). Transcriptomes analysis showed that lung ECs from CKO mice showed an increase of multiple paracrine factors including Cxcl12, Edn1, Pdgfb, and Pdgfd (**Figure 6G**), suggesting that SOX17 deficiency in ECs induces paracrine effect on PASMCs.

**Figure 6.**
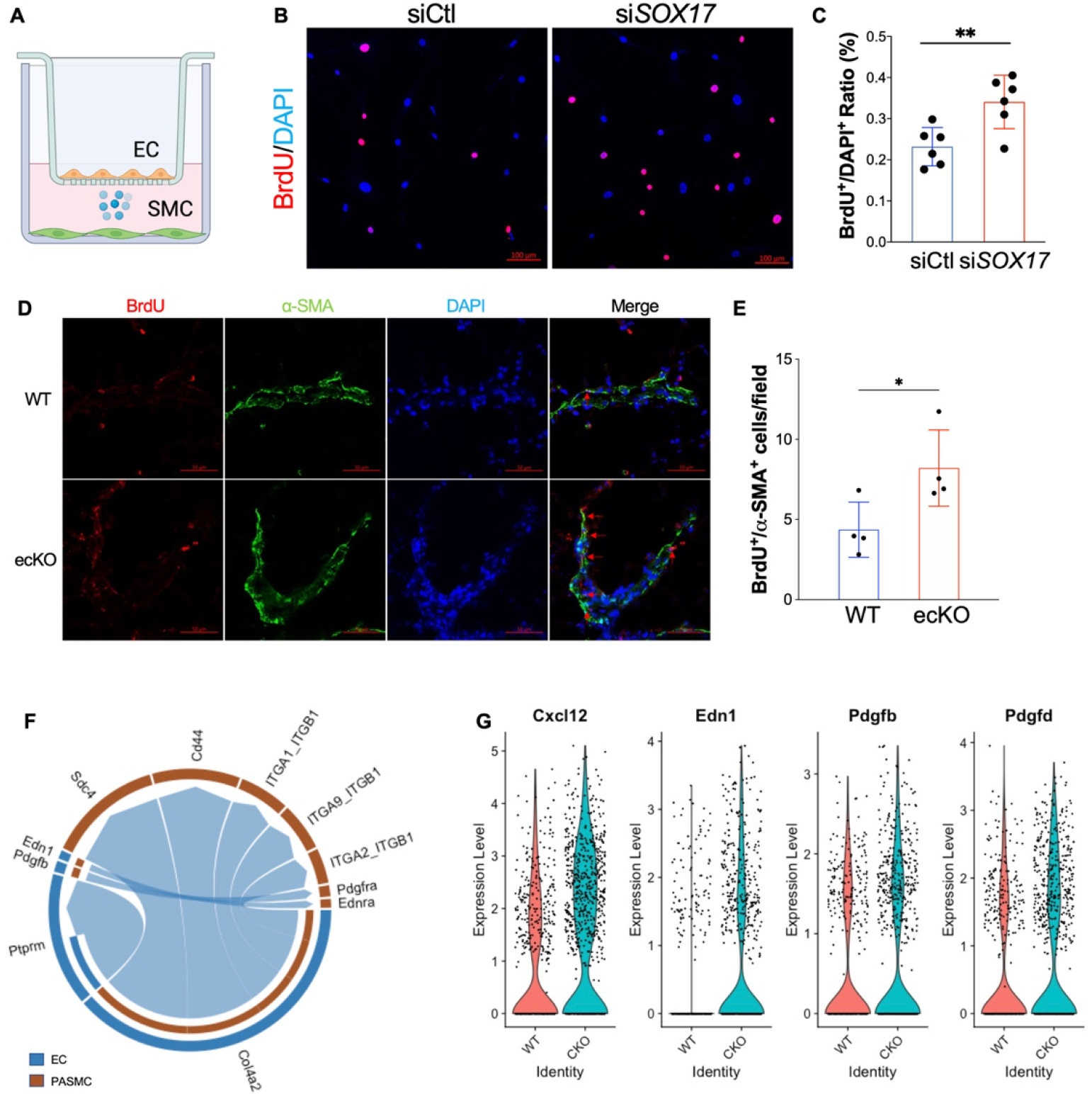
SOX17 deficiency induced PASMC proliferation. (A) A diagram showing the EC and SMCs co-culture model. (B and C) SOX17 deficiency in lung ECs promoted PASMCs proliferation assessed by Transwell co-culture and BrdU assay. PASMCs were seeded on the cover slides on the lower chamber. SOX17 deficiency or control HPVECs were seeded on the top chamber for 48 hours. PASMCs were starved overnight, then co-cultured with HPVECs. BrdU was added in the lower chamber at 8 hours prior to cells harvest. BrdU was stained with anti-BrdU antibodies. Red indicated BrdU positive cells. Nucleus were co-stained with DAPI. (D and E) In vivo BrdU incorporation assay showed upregulation of PASMCs proliferation in ecKO Sox17 mice during hypoxia condition. WT and ecKO Sox17 mice were incubated in hypoxia (10% O_2_) for 10 days. BrdU (25 mg/kg) was injected i.p. between day 7 to day 9. Lung sections were stained with anti-BrdU and anti-*α*-SMA. BrdU^+^/ *α*-SMA^+^ cells were quantified. (F) CellChat prediction using scRNA-seq dataset showed the upregulation of ligand and receptor pairs (Pdgfb-Pdgfra, Edn1-Ednra) in CKO mice. (G) ScRNA-seq analysis showed the increase of EC derived cytokines including Cxcl12, Edn1, Pdgfb, Pdgfd. Student t test (C and E). *, P< 0.05; **, P< 0.01. Scale bar, 50*μ*m.

### SOX17 deficiency induces endothelial dysfunctions

EC hyperproliferation and upregulation of glycolysis are hallmarks of PAH EC (24, 25), we then measure the Extracellular Acidification Rate (ECAR) level and found that SOX17 deficient HPVECs enhanced glycolysis compared to control. (**Figure 7A**). Since anti-apoptotic and hyperproliferative features are hallmarks of PAH ECs, we also evaluated cell apoptosis after SOX17 knockdown. After starvation for 24 hours, HPVECs with SOX17 knockdown exhibited a significant reduction in Caspase 3/7 activity and cleaved Caspase 3 expression, suggesting that SOX17 deficiency promotes anti-apoptotic phenotype of HPVECs **(Figures 7B** and **7C**). Endothelial junction integrity is important to maintain vascular homeostasis. We then measured the EC junction via ECIS system in the presence of Thrombin. Junction integrity is significantly impaired in SOX17 deficient ECs (**Figure 7D**). BMPR2 deficiency is evident in patients with PAH. Our data also demonstrated that SOX17 knockdown reduced BMPR2 expression and BMP9-induced phosphorylation of Smad1/5/9 (**Figure 7E**). These data suggest that SOX17 deficiency induces EC dysfunction including hyperproliferation, enhanced paracrine effect and glycolysis, anti-apoptosis, and impaired junction integrity and BMPR2 signaling leading to EC dysfunction.

**Figure 7.**
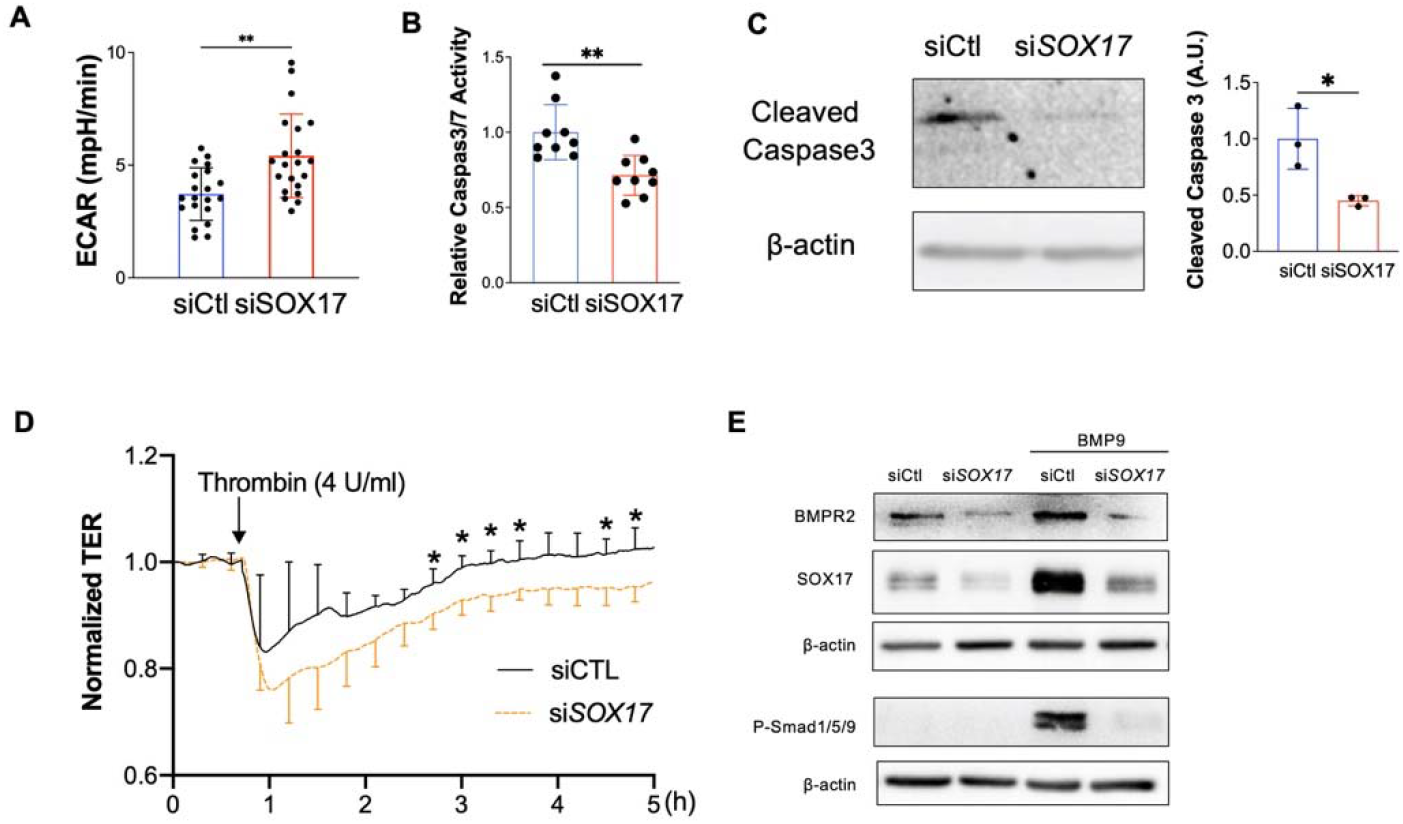
Loss of endothelial SOX17 promoted EC dysfunction. (A) Seahorse glycolytic assay showed that upregulation of Extracellular Acidification Rate (ECAR) levels in SOX17 deficient HPVECs compared to control cells. (B) SOX17 deficiency promoted anti-apoptotic phenotype of HPVECs during starvation assessed by Caspase 3/7 activities. (C) Western blotting analysis demonstrated reduction of cleaved Caspase 3 in SOX17 deficient HPVECs. (D) Impairment of endothelial barrier function in SOX17 deficient HPVECs. At 60 hours post-transfection, TER was monitored for up to 5 hours. Thrombin (4U/ml) was added to disrupt the cellular junction. (n=4). (E) Sox17 deficiency reduced BMPR2 expression and impaired BMPR2 activity via assessing P-Smad1/5/9 expression. Student t test (A-D). *, P< 0.05; **, P< 0.01.

### E2F1 mediated SOX17 deficiency-induced EC dysfunction

To further determine what regulators or transcriptional factors that mediate the upregulation of the proliferative gene program induced by loss of SOX17, we performed transcription factor prediction using iRegulon(26). iRegulon prediction showed that E2F family member *E2F1* is the top transcription factor governing the proliferative program induced by SOX17 deficiency (**Figure 8A**). Western blotting analysis confirmed that SOX17 knockdown markedly induced E2F1 expression in HPVECs (**Figure 8B**). We also observed that E2F1 was significantly upregulated in the lung of ecKO *Sox17* mice (**Figure 8C**). To determine whether E2F1 activation mediates the effect of SOX17 deficiency-induced HPVECs proliferation and survival, we performed siRNA-mediated knockdown of E2F1 in SOX17 deficient HPVECs. siRNA against E2F1 significantly reduced E2F1 mRNA and protein expression (**Figures 8D** and **8E**). We found that E2F1 inhibition via siRNA blocked the expression of cell proliferation genes including *PLK1, CCNB1*, and *CCNB2*, as well as HPVECs proliferation assessed by BrdU incorporation assay (**Figures 8F** and **8G**). Finally, E2F1 knockdown significantly inhibited SOX17 deficiency-induced cell survival (**Figure. 8H**).

**Figure 8.**
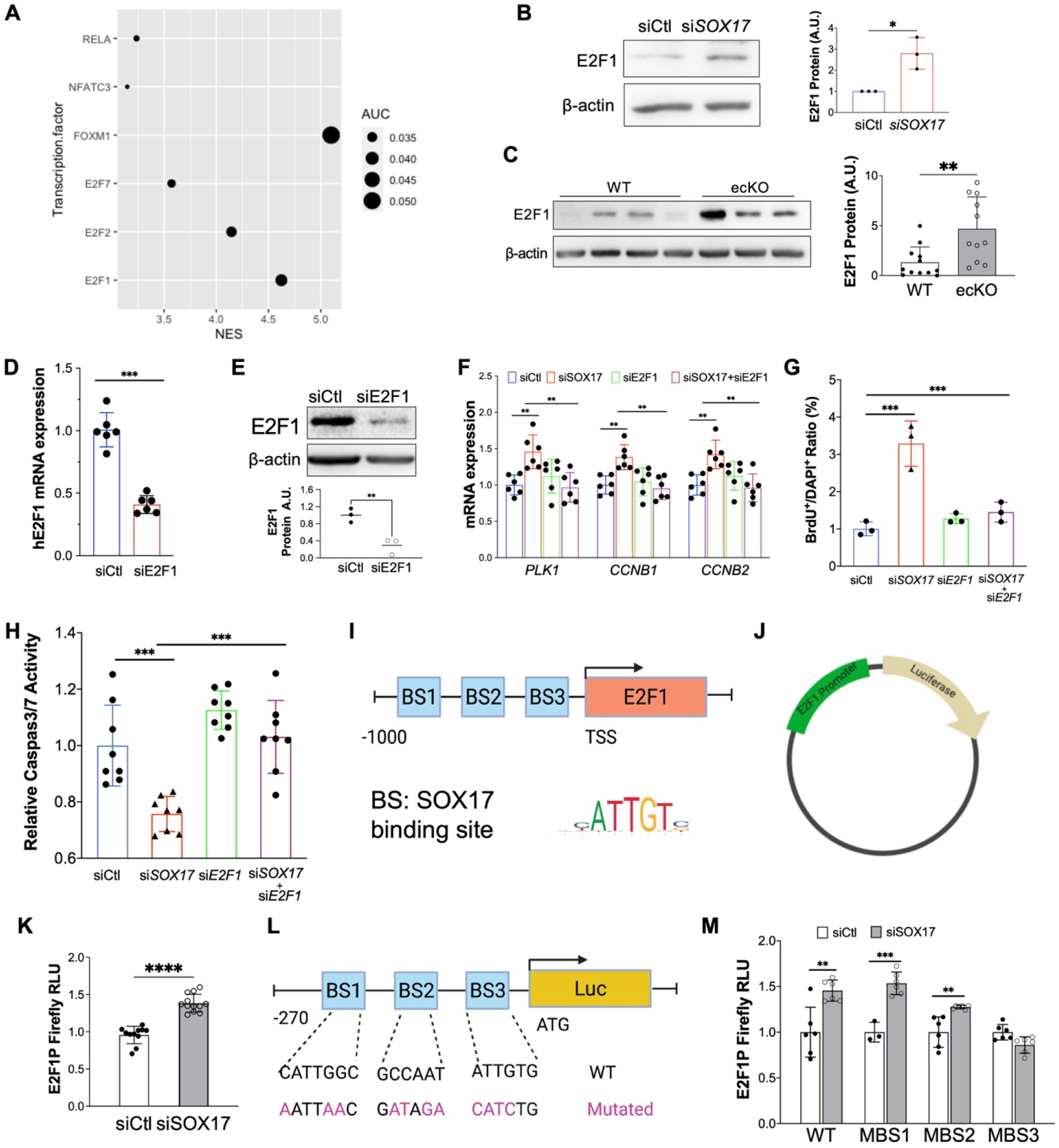
E2F1 mediated SOX17 deficiency-induced dysfunction. (A) iRegulon analysis demonstrated that FOXM1 and E2F1 are the top enriched transcriptional factors potentially governing cell cycle programming in SOX17 deficient HPVECs. (B) Upregulation of E2F1 protein expression by SOX17 knockdown. (C) Increased of E2F1 expression in the lung of ecKO Sox17 mice compared to WT mice. (D) qRT-PCR analysis showed that E2F1 siRNA markedly reduced E2F1 mRNA expression. (E) Western blotting analysis demonstrated that E2F1 protein was efficiently reduced by E2F1 siRNA compared to scramble siRNA. (F) QRT-PCR analysis demonstrated that E2F1 knockdown blocked the genes associated with proliferation including *PLK1, CCNB1, and CCNB2* in the presence of SOX17 deficiency. (G) BrdU incorporation assay demonstrated that E2F1 knockdown normalized cell proliferation induced by loss of SOX17. (H) E2F1 knockdown restored EC apoptosis which was inhibited by SOX17 deficiency. Studies were repeated at least 3 times (B, D, F, G, H). Student t test (C, D and E). (I) A diagram shows that there are 3 putative SOX17 binding sites in the proximal promoter region of human E2F1 gene. (J) A representative map for pLV-E2F1P/Luc plasmid. (K) Loss of SOX17 increased E2F1 promoter activities assessed by luciferase assay. HPVECs were transfected with control of SOX17 siRNA for 12 hours, followed by infected with pLV-E2F1P/luc lentivirus for 48 hours. (L) A diagram showing that the SOX17 putative binding sites in E2F1 promoter/luciferase constructs were mutated. Purple highlight letters indicate mutated DNA sequences of the SOX17 putative binding sites in the E2F1 promoter. (M) Binding site 3 mutation blocked SOX17 deficiency-induced E2F1 promoter activation. MBS1/2/3 indicate mutated binding site 1/2/3. HPVECs were transfected with control of SOX17 siRNA for 12 hours, followed by infected with WT or mutated pLV-E2F1P/luc lentiviruses for 48 hours. Studies were repeated at least 3 times. One-way ANOVA with Tukey post hoc analysis (F, G and H). Student t test (K and M). *, P< 0.05; **, P< 0.01, ***, P< 0.001, ****, P< 0.0001.

### Transcriptional upregulation of E2F1 promoter is activated by SOX17 deficiency

To characterize whether *E2F1* is a direct transcriptional binding target of SOX17 in HPVECs, we did *in silico* promoter analysis (Eukaryotic Promoter Database)(27) of the human *E2F1* promoter and found that there are 3 putative SOX17 binding sites in the human *E2F1* proximal promoter (−200bp to +1bp of TSS) (**Figure 8I**). We then cloned the E2F1 promoter into the upstream of luciferase gene (**Figure 8J**). Knockdown of SOX17 significantly upregulated the promoter activity of E2F1 assessed by luciferase assay (**Figures 8K**), suggesting that SOX17 might repress E2F1 through binding to SOX17 binding sites in the promoter of E2F1. To determine which putative binding sites in the E2F1 promoter are response for E2F1 suppression by SOX17, we mutated individual binding site and co-transfected with SOX17 siRNA (**Figure 8L**). Our data showed that binding site 3 mutation inhibited SOX17 deficiency induced E2F1 promoter activation, suggesting that binding side 3 is likely the binding region of SOX17 in E2F1 promoter in lung ECs (**Figures 8M**).

### E2F1 signaling inhibition rescued SOX17 deficiency-induced PH in mice

To determine whether E2F1 is involved in SOX17 deficiency-induced EC dysfunction, E2F1 inhibitor (HLM) was added to in HPVECs for 6 hours. BrdU assay and qRT-PCR and Western Blot analysis showed that E2F1 inhibition significantly impeded cell proliferation and the levels of the genes (*PLK1, CDKN2C, CCNA2*) related to cell proliferation. (**Figures 9A-9C**). We also found that E2F1 inhibition rescued the anti-apoptotic phenotype and paracrine effect of SOX17 deficient HPVECs. (**Figure 9D and 9E**). To further determine the therapeutic potential of targeting E2F1 signaling, we treated ecKO *Sox17* mice with HLM or vehicle (**Figure 9F**). We found that HLM treatment almost completely rescued the PH phenotype, as RVSP levels was significantly reduced by HLM treatment compared to vehicle (**Figure 9G**). The RV/LV+S ratio was not changed by the treatment of HLM (**Figure 9H)**. Further examination of pulmonary pathology showed that the muscularization of distal pulmonary arteries and pulmonary wall thickness were markedly attenuated by HLM treatment (**Figures 9I-9L**). Collectively, our studies suggest that E2F1 signaling mediates SOX17 deficiency-induced PH in mice and targeting E2F1 represents a novel therapeutic approach for the treatment of PH with SOX17 deficiency.

**Figure 9.**
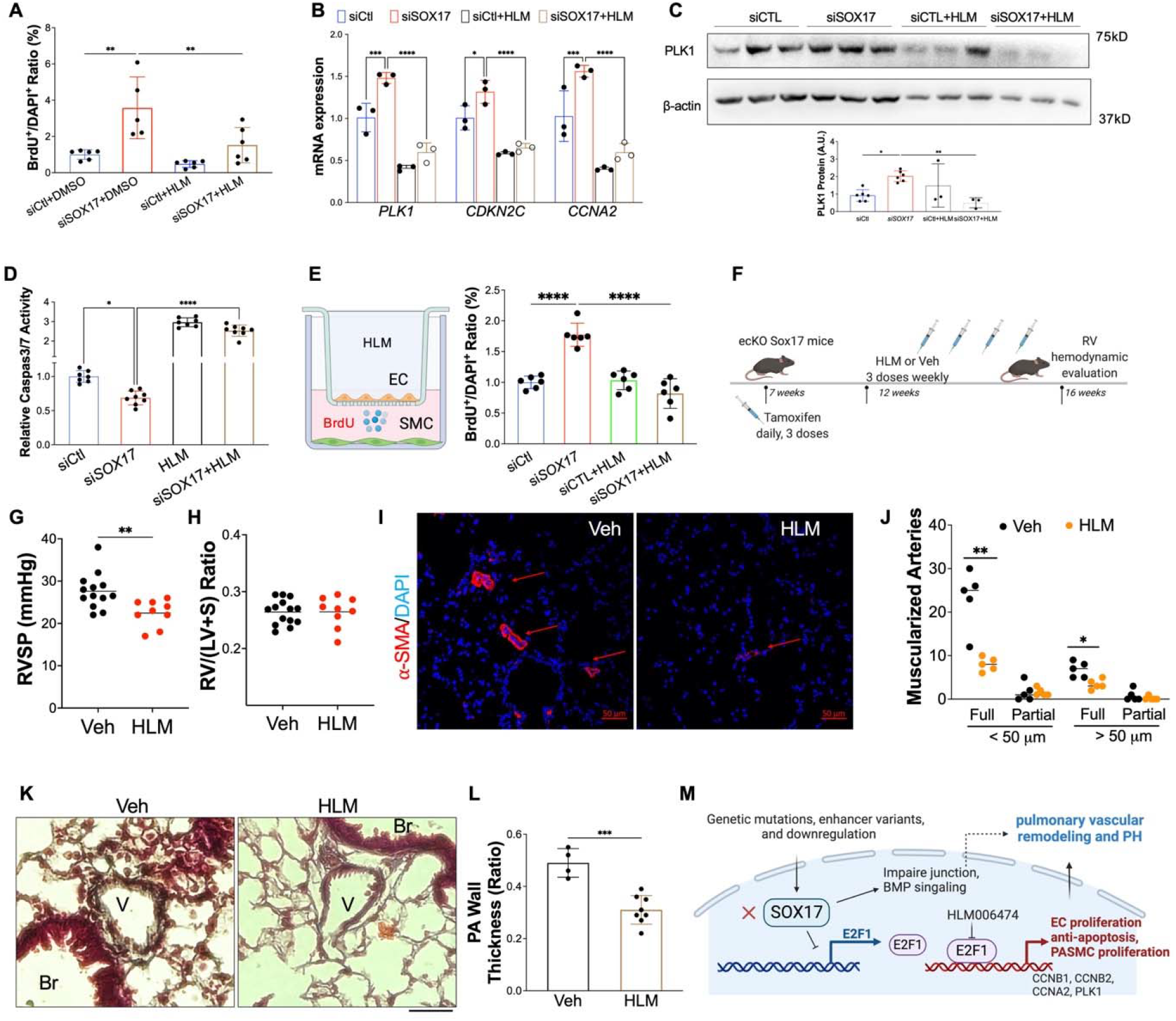
Pharmacological inhibition of E2F1 reduced EC dysfunction and PH development in ecKO *Sox17* mice. **(A)** E2F1 inhibition reduced EC proliferation measured by BrdU incorporation assay. At 48 hours post-transfection of siRNA against SOX17 or control siRNA, HPVECs were treated with DMSO or HLM for 12 hours in serum/growth factors free medium. 2.5% FBS and BrdU were added in the medium at 4 hours prior to cells harvest. (B) qRT-PCR analysis demonstrated normalization of the expression of genes related to cell proliferation after E2F1 inhibition in HPVECs. At 48 hours post-transfection, HPVECs were treated with DMSO or HLM for 12 hours in serum/growth factors free medium. 2.5% FBS were added in the medium at 4 hours prior to RNA isolation. (C) E2F1 inhibition reduced cell proliferation marker PLK1 expression in SOX17 deficiency in HPVECs. (D) Pharmacological inhibition of E2F1 increased EC apoptosis in SOX17 deficient HPVECs. At 48 hours post-transfection, HPVECs were treated with DMSO or HLM for 12 hours in serum/growth factors free medium, followed by measurement of Caspase 3/7 activities. (E) A diagram showing the strategy of E2F1 inhibition in ecKO Sox17 mice. (F) RVSP was attenuated by E2F1 inhibition in ecKO Sox17 mice. (G) RV hypertrophy was not altered by E2F1 inhibition. (H and I) Muscularization of distal pulmonary arteries were reduced by E2F1 inhibition in ecKO Sox17 mice compared to vehicle. *α*-SMA^+^ vessels were quantified in 20 field at 10X magnification per mouse. (J and K) Pentachrome staining showed that E2F1 inhibition by HLM attenuated pulmonary wall thickness. Wall thickness was calculated by the distance between internal wall and external wall divided by the distance between external wall and the center of lumen. Studies were repeated at least 3 times (A, B, D). One-way ANOVA with Tukey post hoc analysis (A-D) and Student t test (F, G, I, K). *, P< 0.05; **, P< 0.01, ***, P< 0.001, ****, P< 0.0001. Scale bar, 50*μ*m.

## Discussion

The present study has demonstrated that genetic disruption of *Sox17* in ECs induces mild PH as evident by increased RVSP and pulmonary vascular remodeling. We also observed that SOX17 expression is significantly downregulated in isolated PAECs from IPAH patients and is diminished in the occlusive vessels of IPAH lungs. In addition, we found the increased cell proliferation, survival and paracrine effect, impairment of cellular junction and BMP signaling in SOX17 deficient PAECs. We then demonstrated that E2F1 is induced by loss of SOX17 and mediates the cell dysfunctions induced by SOX17 deficiency. Pharmacological inhibition of E2F1 attenuated PH in ecKO *Sox17* mice. These findings raise the exciting possibility that inhibition of E2F1 signaling could treat PAH patients with SOX17 deficiency (**Figure 9M**).

Endothelial dysfunction is believed to be the initial event during the development of PAH(28). Single-cell transcriptomics analysis showed that expression of SOX17 is preferentially expressed in the lung ECs compared to other cell types. However, SOX17 expression is markedly downregulated in the lung ECs isolated from IPAH patients and the lung of MCT-induced PH models, suggesting that EC SOX17 deficiency mediates the development of PAH in patients.

Endothelial dysfunctions including hyperproliferation and anti-apoptosis are hallmark of PAH (24, 25, 29). Increased cell proliferation and apoptosis-resistance were evident in the SOX17-deficienct ECs. SOX17 deficiency also led to upregulation of glycolysis, one of important mechanisms mediating EC dysfunction in PAH(30). We also observed that loss of SOX17 resulted in impairment of cellular junction integrity and BMP signaling, important features of lung vasculature in maintaining lung hemostasis. Loss of SOX17 in ECs also enhanced the paracrine effect such as promotion of PASMCs proliferation. Our scRNA-seq analysis also indicated there might be deficiency of lung arterial EC differentiation in Sox17 deficiency lung (**Supplemental Figure 5**), as SOX17 is critical for maintaining arterial identity(8), which is consistence with recent study showing Notch1 deficiency due to Sox17 loss in mice(16). Using tamoxifen-inducible EC-specific Sox17 deletion in adult mice, our work demonstrated the causal role of SOX17 deficiency in inducing endothelial dysfunction, pulmonary vascular remodeling and the development of PH. This observation is consistent with the finding that SOX17 mutations were present in patients with IPAH and congenital heart disease associated PAH(13, 14).

In addition to PAH patients, SOX17 expression is downregulated in many forms of cancer, including colorectal cancer(31), breast cancer(32), endometrial cancer(33), and cholangiocarcinoma(34), due to DNA hypermethylation at SOX17 promoter loci. Mechanistically, SOX17 serves as a tumor suppressor through the suppression of tumor cell proliferation and migration via modulation of Wnt signaling(34–36). Reduced SOX17 expression is also present in the intracerebral arteries of intracerebral aneurysm patients(37). Deficiency of SOX17 in ECs induces intracerebral aneurysm (37). Other studies demonstrated that EC-specific inactivation of *Sox17* in mice leads to brain microcirculation leakage due to loss of Wnt/β-catenin signaling(38). It seems that β-catenin is not involved in the pro-proliferation and anti-apoptosis phenotypes of SOX17 deficient HPVECs, as β-catenin knockdown did not block the pro-proliferation effect induced by loss of SOX17 (**Supplemental Figure 6**).

Using unbiased analysis of the single-cell and bulk transcriptomes altered by SOX17 deficiency, we identified cell proliferation and paracrine effect program (including Pdgfb, Edn1, Cxcl12) is upregulated by loss of SOX17 in vitro and in vivo. Other study also showed that combined Sox17 deficiency with hypoxia induced HGF signaling and endothelial proliferation in vivo(16). We then predicted and validated that E2F1 is the central governor controlling the EC dysfunction by SOX17 deficiency. E2F1 belongs to a subclass of the E2F transcription factor family and is thought to act as a transcriptional activator, mediating cell proliferation and apoptosis(39, 40). E2F1 is critical for the expression of various genes regulating G1 to S transition and S phase, including cyclin E, PCNA, Ki67, BUB1, Cyclin A2, Cyclin B1, Cyclin B2, etc(41, 42). Loss of E2F1 was shown to mediate TNF-α-induced cell cycle arrest in proliferating bovine aortic ECs(43). Restoration of E2F activities via adenovirus-mediated E2F1 overexpression promoted EC cell cycle progress and rescued TNF-α-induced apoptosis(43). Our studies demonstrated that E2F1 expression and promoter activities are upregulated by SOX17 deficiency in HPVECs likely due to absence of suppression of SOX17 in the proximal region of E2F1 promoter. Moreover, E2F1 has been shown to mediate sodium–hydrogen exchanger 1 (NHE1) induced PASMCs proliferation, hypertrophy and migration in vitro(44). E2F1 expression is also significantly increased in the lung of other PH models such as monocrotaline-exposed rats(45) and *Egln1^Tie2Cre^* mice(17, 46) (**Supplemental Figures 7A and 7B**). Overexpression of E2F1 suppressed BMPR2 expression in the HPVECs (**Supplemental Figure 7C**). Taken together, E2F1 activation is likely the common mechanisms mediating pulmonary vascular remodeling and PH development.

The present study has demonstrated that targeting E2F1 signaling with HLM effectively inhibited Sox17 deficiency-induced PH development in mice. Pharmacological inhibition of E2F1 reduced HPVECs pro-proliferation, anti-apoptotic phenotypes and paracrine effect due to SOX17 deficiency and pulmonary vascular remodeling and PH in ecKO Sox17 mice. It is possible that E2F1 inhibition also reduced PASMCs proliferation in ecKO mice. Other studies showed that inhibition of E2F1 signaling prevented occlusive thickening of the vessel wall in venous bypass grafts(47). Future studies are warranted to investigate whether or not E2F1 inhibition could attenuate PH development and right heart dysfunction in more severe PH models such as MCT-exposed rat, SuHx-rats, or *Egln1^Tie2Cre^* mice.

In summary, our studies demonstrate a pathogenic role of endothelial SOX17 deficiency in mediating lung EC proliferation/anti-apoptosis and pulmonary vascular remodeling, and provide clear evidence of E2F1 activation in the pathogenesis of PH. We also show that pharmacologic inhibition of E2F1 attenuated PH development in ecKO Sox17 mice. These studies suggest that E2F1 inhibition could be a promising approach for the treatment of PAH patients with loss of SOX17 or E2F1 activation.

## Author contributions

Z.D. conceived the experiments and interpreted the data. D.Y., B.L., R.L., J. P., I. S., R. F., W.H.L, Y. Z, and Z.D. designed, performed experiments, and analyzed the data. Z.D. wrote the manuscript. W.H.L., T.W., C.C.G, revised the manuscript. M.B.F. provided key experimental materials.

## Acknowledgements

We thank Dr. Marlene Rabinovitch (Stanford University) for her advice on the experimental design and data interpretation. The authors the Pulmonary Hypertension Breakthrough Initiative for providing the Data/tissue samples. Funding for the Pulmonary Hypertension Breakthrough Initiative is provided under an NHLBI R24 grant (R24HL123767) and by the Cardiovascular Medical Research and Education Fund.

