## Supplemental materials for "E2F1 Mediates SOX17 Deficiency-Induced Pulmonary Hypertension"

**Materials and Methods**

**Human samples**

The use of archived human lung tissues and cells were granted by the University of Arizona (UA) Institutional Review Board**.** Human IPAH patients and failed donors (FD)’ pulmonary microvascular ECs (PMVECs) were obtained from the Pulmonary Hypertension Breakthrough Initiative (PHBI). Both IPAH and FD PMVECs were isolated from arteries that are less than 1mm thickness according to cell isolation protocol provided by PHBI. A table summarizing clinical and demographic characteristics of IPAH patients and FD are provided in **Supplemental Table 1**. The normal human pulmonary ECs (HPVECs) were purchased from Lonza (Alpharetta, GA, USA)

**Mice**

Sox17 floxed (*Sox17*^f/f^) mice (JAX#031712) were purchased from Jackson Laboratory. cKO Sox17 mice were generated by breeding *Sox17*^f/f^ mice with Tie2Cre mice(1). ecKO *Sox17* mice were generated by breeding *Sox17*^f/f^ mice with *EndoSCL-CreERT2* mice(2). Both male and female mice were included for experiments. At age of 7 weeks old, ecKO *Sox17* mice and their *Sox17^f/f^* (WT) littermates were treated with tamoxifen (Sigma, #T5678) at the dose of 100 mg/kg body weight (i.p., daily) for 3 continue days to induce genetic deletion. For normoxic experiment, two months post tamoxifen treatment, mice were used for hemodynamic measurement and tissue collection. For hypoxia studies, 3 weeks post tamoxifen treatment, mice were incubated in 10% O_2_ hypoxia chamber (Biosphere) for three weeks, followed by hemodynamic measurement and tissue collection. For HLM treatment, ecKO *Sox17* mice were treated with tamoxifen as described above. Two weeks post treatment, mice were treated with HLM006474 (HLM, 12.5 mg/kg) 3 times a week for 6 weeks, followed by hemodynamics measurement and tissue harvest. The studies were conducted according to National Institutes of Health guidelines on the use of laboratory animals. The protocol for animal care and studies was approved by the Institutional Animal Care and Use Committee of UA.

**Hemodynamic and echocardiography measurement**

Right ventricular systolic pressure (RVSP) was measured with a 1.4F pressure transducer catheter (Millar Instruments) and recorded with AcqKnowledge software (Biopac Systems Inc.) as described previously(1, 3). Briefly, the catheter was inserted into the right ventricle in mice under anesthesia (100 mg ketamine/ 5mg xylazine/ kg body weight, i.p.).

Echocardiography was performed at the Translational Cardiovascular Research Center at University of Arizona. Transthoracic echocardiography was performed on a VisualSonics Vevo 3100 ultrasound machine (FujiFilm VisualSonics Inc) using an MS550D (40 MHz) transducer. The left ventricle size and function including fractional shortening (LV FS), the cardiac output (CO), and heart rate were obtained from the parasternal short axis view using M-mode. The RV cross-sectional area was obtained from the parasternal short axis view at the papillary muscle level using B-mode. Pulmonary artery (PA) acceleration time and PA ejection time were obtained from the parasternal short axis view at the aortic valve level using pulsed Doppler mode.

**Immunofluorescent staining and histological staining**

Human IPAH patients and failed donors’ lung sections were fixed with 4% paraformaldehyde for 20 min. The sections were blocked with 5% goat serum and incubated with anti-SOX17 (1:1000, Abcam, Cat#ab224637) and Alexa Fluor 647 conjugated anti-CD31 (1:20, Santa Cruz Biotechnology, Cat#sc-376764 AF647) were incubated at 4 °C overnight, followed by 1 hour incubation with Alexa 594 conjugated anti-rabbit IgG (Thermo Fisher Scientific, Waltham, MA, USA) at room temperature for 1 h. Nuclei were counterstained with mounting media containing DAPI (SouthernBiotech, Birmingham, AL, USA).

For immunofluorescent staining, the left lobe of lung tissue was perfused with 50% OCT in PBS through tracheal catheterization, and then embedded in OCT for cryosectioning. Lung sections (5 μm) were fixed with 4% paraformaldehyde, followed by blocking with 0.1% Triton X-100 and 5% normal goat serum at room temperature for 1 h. After 3 washes, they were incubated with anti-α-smooth muscle actin (α-SMA, 1:300, Abcam, Cat#ab5694) or anti-SOX17 (1:1000, Abcam, Cat#ab224637) or anti-CD31 (1:25, BD Bioscience, Cat#550274) at 4 °C overnight and were then incubated with fluorescence-conjugated secondary antibodies (Life Technology) at room temperature for 1 hour after 3 washes. Nuclei were counterstained mounting media containing DAPI (SouthernBiotech, Birmingham, AL, USA).

For immunohistochemistry and histology, well-perfused lungs were fixed for 5 min by instillation of 10% PBS-buffered formalin through trachea catheterization at a transpulmonary pressure of 15 cm H_2_O, and then 48 h fixation at room temperature on shaker. After paraffin processing, the tissues were cut into semi-thin 5 μm thick sections. Mouse lung sections were then dewaxed and dehydrated. Antigen retrieval was performed by boiling the slides in antigen retrieval buffer (Vector Lab) for 1 minutes in high pressure cooker. After blocking, slides were incubated with anti-α-SMA (1:300, Abcam, Cat#ab5694) at 4 °C overnight and were then incubated with Alexa 594 conjugated anti-rabbit IgG at room temperature for 1 hour. The nuclei were counterstained with DAPI. α-SMA^+^ vessels were quantified in 20 field at 10X magnification per sample.

For assessment of vascular remodeling, paraffin sections of lungs were dewaxed and dehydrated, and then stained with a Russel-Movat pentachrome staining kit (Statlab) as described previously(3, 4). For assessment of PA wall thickness, PAs from images at 20X magnification were quantified by Image J. Wall thickness was calculated by the distance between internal wall and external wall divided by the distance between external wall and the center of lumen as described previously(5).

**Primary cultures of human lung vascular ECs.**

Primary HPVECs were purchased from Lonza (#CC-2527). Briefly, HPVECs were cultured in 0.2% gelatin (Sigma) precoated T75 flasks in EBM-2 medium supplemented with 10% FBS and EGM-2 MV singlequots (Lonza, #CC-3162), and incubated at 37 °C in a humidified 5% CO_2_ atmosphere. Cells between passage 4 and 7 were used for experiments. Primary IPAH patients and FD PVECs were obtained from PHBI and cultured as described above.

**siRNA-mediated gene knockdown**

SOX17 (#sc-38429), E2F1 (#sc-29297) and control siRNA (#sc-37007) were purchased from Santa Cruz Biotechnology. siRNA (50μM) was transfected into ECs using HMVEC-L Nucleofector kit (Lonza, # VPB-1003) with Nucleofector^TM^ 2b device (Lonza, #AAB-1001) with program A-034. For BMP9 treatment, HPVECs were knocked down with SOX17 and control siRNA for 48 hours, followed by BMP9 (5 ng/ml, BioLegend, Cat#553102) incubation for 16 hours.

***In vitro* cell proliferation and apoptosis analysis**

For assessment of HPVECs proliferation after SOX17 or E2F1 knockdown, HPVECs were transfected with SOX17 siRNA or scramble siRNA and seeded on coverslips to reach 40% to 60% confluency. Around 48 hrs post transfection, cells were starved overnight in FBS-free medium. 2.5% FBS and 5-bromo-20-deoxyuridine (BrdU, Sigma, Cat #B5002) were added (10 µM) for another 4 hrs. For inhibitors studies, E2F1 inhibitor HLM006474 (HLM, 40 μM) were administrated 2 hr before FBS and BrdU treatment. Fixed cells were immunostained with anti-BrdU antibody (BD bioscience, cat#347580), followed by Alexa 594-conjugated anti-mouse IgG (Life Technology). Nuclei were counterstained with DAPI. Images were taken using REVOLVE microscopy (Discover Echo Inc.) at 4 X magnification for quantification. For assessment of EC apoptosis, HPVECs after transfection were seeded on 96 well plate for 48 hrs. Cells were starved for overnight and used for Caspase 3/7 activity assay for apoptosis detection. (Promega, #G8090). For inhibitor studies, HLM (40 μM) was added 6 hours before sample harvest.

**RNA sequencing and bioinformatic analysis**

HPVECs were transfected with control siRNA (siCtl) or SOX17 siRNA for 48 hours. Total RNA was isolated from cells with Zymo Research Quick-RNA Miniprep Kits with DNase Ι digestion. Three replicates were pooled in the equal amount for RNA-seq. RNA sequencing was carried out at the Novogene Corporation Inc. All samples were sequenced with an Illumina Novaseq 6000 platform with a pair end 150-bp read length. For bioinformatics analysis, raw reads were aligned with Hisat2 against the reference genome plus the transcriptome sequence by using UCSC Refseq annotations. Per-gene expression was quantified and the differential expression, fold-changes between conditions using CuffDiff. The RNA sequencing raw data and analyzed data are available at the National Center for Biotechnology Information Gene Expression Omnibus Database (GSE192649, the data will be released after being published). For pathway enrichment analysis, genes with FKPM > 5 and fold change > 1.5 were used as an input in DAVID Bioinformatics Resources 6.8. For transcription factor prediction, the filtered genes were used an input and analyzed via iRegulon plugin in Cytoscape 3.2 software.

SOX17 binding sites prediction of the human E2F1 promoter was performed *in silico* through Eukaryotic Promoter Database(6). The promoter regions spanning 1000bp upstream and 100 downstream to TSS were used for analysis. The cut-off P value is set at 0.001.

**Reanalysis of public microarray and single-cell RNA sequencing datasets**

The public microarray dataset (GSE113439) for IPAH patients were analyzed with GEO2R provided by the NCBI website(7). We also used the publicly available single-cell RNA sequencing metadata from human lung (GSM3926539)(8). The metadata was processed in R (version 4.0.2) via Seurat package V4. Briefly, cells that expressed fewer than 100 genes, and cells that expressed over 4,000 genes, and cells with unique molecular identifiers (UMIs) more than 10% from the mitochondrial genome were filtered out. The data were normalized and integrated in Seurat, followed by Scaled and summarized by principal component analysis (PCA), and then visualized using UMAP plot. FindClusters function (resolution = 0.3) in Seurat was used to cluster cells based on the gene expression profile. Endothelial cells (expressing CDH5), fibroblasts (FN1, COL1A), smooth muscle cells (ACTA2), type 1 alveolar epithelial cells (SFTPB), type 2 alveolar epithelial cells (SFTPC), club ciliated cells (SCGB1A1), pericytes (PDGFRB), Mast cells (MCTP1), PMN (S100A9) and macrophages (CD68, CD14), T cells (CD3G), dendric cells (IRF8, MS4FA1) clusters were annotated based on the expression of known markers.

**single-cell RNA sequencing analysis**

We collected whole lung tissues from WT and cKO mice at the age of 2 months for scRNA-seq analysis. Equal number of single cells isolated from WT and CKO mice was stained with Cell Hashing antibodies (Biolegend, Cat#155831 and #155833), respectively, and loaded on 10X Genomics Chromium Single Cell Controller to generate barcoded single cells for construction of single cell cDNA libraries. ScRNA-seq was performed on a Hiseq 4000 with pair-end 150bp. ScRNA-seq data were prefiltered to remove cells with negative hashing tag or doublets and high mitochondrial transcripts (> 15%) using Seurat v4(9). Employing Seurat v4 for graph-based clustering analysis(9), we revealed discrete cell populations in mouse lungs, including ECs (*Pecam1*, *Cdh5*), SMCs (*Acta2*, *Myh11*), fibroblasts (*Col1a1* and *Fn1*), pericytes (*Pdgfrb*, *Notch3 and Cspg4*), alveolar type 1 (*Aqp5*, *Ager*) and type 2 (*Sftpb*, *Sftpc* and *Abca3*) epithelial cells, macrophages (*Cd68*, *Cd14*, *Spi1* and *Fcgr2b*), T lymphocytes (*Cd3g*, *Cd3e* and *Cd7*), B lymphocytes (*Ighm*), neutrophiles (PMNs) (*S100a9, S100a8*), natural killer cells (NK) (Klrc1) predicted by cell type-selective markers(10) using UCell package(11). Lung ECs transcriptomes were extracted for differential gene expression, EC subpopulation clusters, and CellChat prediction(12). For lung EC subpopulations clustering, we defined EC subpopulation according to their markers(10, 13). To predict the ligand-receptor interaction between ECs and SMCs, CellChat pipeline was used(12).

**QRT-PCR analysis**

Total RNA was isolated from human PVECs or HPVECs using a Zymo Research Quick-RNA Miniprep Kits with DNase Ι digestion. RNA from mouse and human cells were directly isolated with the RNeasy mini kit. One microgram of RNA was transcribed into cDNA using the high- capacity cDNA reverse transcription kits (Applied Biosystems) according to the manufacturer’s protocol. Quantitative RT-PCR analysis was performed on an QuantStudio 3 system (Applied Biosystems) with the Power Up SYBR Green Master kit (Applied Biosystems). Target mRNA was determined using the comparative cycle threshold method of relative quantitation. Cyclophilin was used as an internal control for analysis of expression of mouse genes 3 while 18s rRNA gene was used for human genes. The primer sequences are provided in **Supplemental Table 2.**

**Western Blot**

Protein lysates were prepared by mincing lung tissues from MCT-exposed rats(4) and mice or harvesting cells in cold NP-40 lysis buffer supplemented with protease inhibitor cocktails (MilliporeSigma). Equal amount of protein was loaded for SDS-PAGE and Western Blotting. The PVDF membranes were blotted with anti-SOX17 (Abcam, #ab224637, 1:10,000), anti-E2F1 (Santa Cruz Biotechnology, #sc-251, 1:500), anti-Cleavage Caspase 3 (Cell Signaling Technology, #9661, 1:1000), anti-BMPR2 (1:1000, BD Biosciences, Cat#612292), anti-P-Smad1/5/9 (1:1000, Cell Signaling Technology, Cat#13820S) or anti-β-actin (Sigma-Aldrich, #A2228) antibodies. The quantification of densitometric measurements for blot images was performed on the Image J.

**Transendothelial Monolayer Electrical Resistance (TER) Assay**

Endothelial monolayer junctional changes were measured by an electric cell substrate impedance sensor (ECIS) to record real-time changes in electrical resistance across endothelial monolayers. Briefly, HPVECs after transfection with SOX17 siRNA were seeded at confluence on a ECIS chamber with small gold electrode (Applied Biophysics) overnight. Before the experiment, the confluent endothelial monolayer was monitored for 0.5 hour at baseline, the endothelial monolayers were treated with Thrombin (4U/ml) and monitored up to 12 hours. The data was normalized to the baseline value.

**Luciferase assay**

E2F1 promoter fused luciferase gene plasmids (#246573) was obtained from Addgene. The DNA sequences containing E2F1 promoter (E2F1-P) and luciferase gene (Luc) were cloned into pLV-mCherry vector (Addgene, #36084) via replacing CMV-promoter and mCherry gene with E2F1-P and Luc. Helper plasmids PSPAX2 vector (Addgene, #12260) and pMD2.G vector (Addgene, #12259) were co-transfected with pLV-E2F1-P/Luc plasmids using lipofectamine 3000 (Thermo Fisher) into 293-T cells to generate lentivirus.

HPVECs was transfected with SOX17 or control siRNAs for 12 hours, followed by infection of pLV-E2F1-P/Luc lentivirus for 48 hours. Cells were lysed for Steady-Glo® Luciferase Assay (Promega). Mutations of putative binding sites in the E2P1 promoter were performed via overlapping PCR and ligated with pLV vector.

**Data availability**

RNA-seq and scRNA-seq data have been deposited in the GEO database under accession number GSE192649 and GSE218398. Scripts used for single-cell RNA sequencing analysis and analyzed data in R objects are available in Figshare (https://figshare.com/s/37782988b8cac7cedcf9). Other data that support the findings of this study are available from the corresponding author upon reasonable request.

**Statistical Analysis**

Statistical determination was performed on Prism 9 (Graphpad Software Inc.). Two-group comparisons were compared by the unpaired 2-tailed Student t test for equal variance or the Welch t test for unequal variance. Multiple comparisons were performed by One Way ANOVA with a Tukey post hoc analysis that calculates corrected P values. P less than 0.05 indicated a statistically significant difference. All bars in dot plot figures represent mean. All bar graphs represent mean±SD.

**Supplemental Table 1. Clinical and demographic characteristics of PAH patients and control**

| **PAH or Failed Donor** | **Gender** | **Race** | **Age** |
| --- | --- | --- | --- |
| IPAH1 | Female | White | 11 |
| IPAH2 | Male | White | 53 |
| IPAH3 | Female | White | 55 |
| IPAH4 | Male | White | 40 |
| IPAH5 | Male | White | 51 |
| IPAH6 | Female | White | 16 |
| Failed donor 1 | Female | White | 34 |
| Failed donor 2 | Female | White | 55 |
| Failed donor 3 | Female | White | 49 |
| Failed donor 4 | Female | White | 57 |
| Failed donor 5 | Male | White | 30 |
| Failed donor 6 | Female | White | 46 |
| Failed donor 7 | Male | Unknown | 20 |
| Failed donor 8 | Male | White | 30 |
| Failed donor 9 | Female | White | 50 |
| Failed donor 10 | Male | White | 51 |
| Failed donor 11 | Male | unknown | 57 |

**Supplemental Table 2. qRT-PCR primers**

| Gene name | Forward primer | Reverse primer |
| --- | --- | --- |
| *SOX17* | GAGCCAAGGGCGAGTCCCGTA | CCTTCCACGACTTGCCCAGCAT |
| *CKDN2C* | ACTGCGCTGCAGGTTATGAA | AGCGAAACCAGTTCGGTCTT |
| *CCNB2* | GCACAAGTAGCTAAGAAAGCTCA | CTCAGGTGTGGGAGAAGGAC |
| *CCNB1* | ACCTGTGTCAGGCTTTCTCTG | CTGACTGCTTGCTCTTCCTCA |
| *CCNA2* | GCATGTCACCGTTCCTCCTT | GGGCATCTTCACGCTCTATTT |
| *PLK1* | CACAGTGTCAATGCCTCCAAG | TATCACAGAGCTGATACCCAAG |
| *E2F1* | ACTGACCATCAGTACCTGGC | CGGGGATTTCACACCTTTTCC |
| *18s rRNA* | TTCCGACCATAAACGATGCCGA | GACTTTGGTTTCCCGGAAGCTG |

**Supplemental Figure legends.**

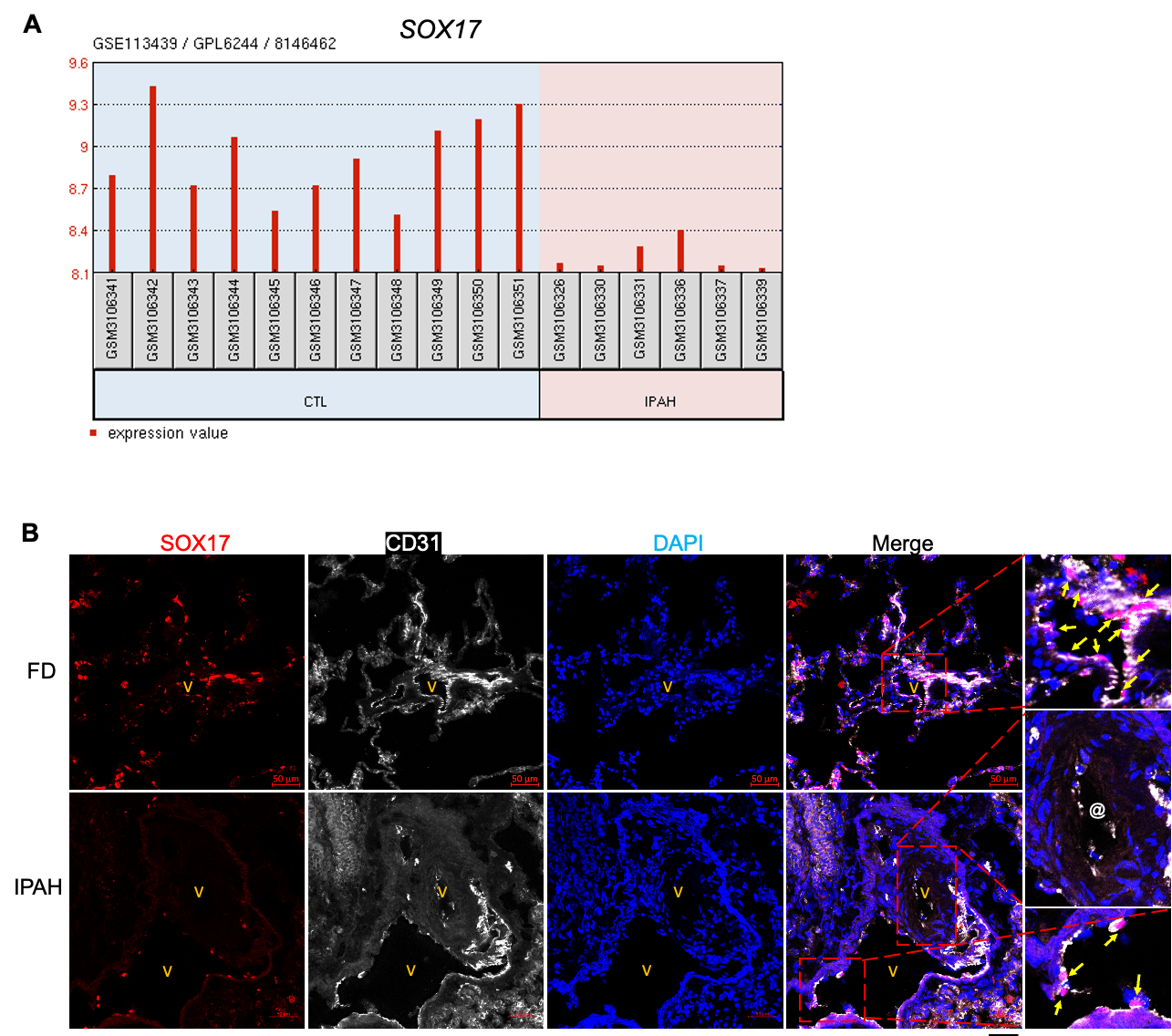

**Supplemental Figure 1. Downregulation of SOX17 in IPAH patients.** (A)Public microarray data showing the downregulation of SOX17 mRNA in the patients with PAH. CTL=non-PAH donors. (B) Immunostaining showed the downregulation of SOX17 in lung ECs from IPAH patients. FD=failed donors. @ indicates diminished SOX17 expression in the occlusive vessels of IPAH patients. Scale bar, 50$\mu$m

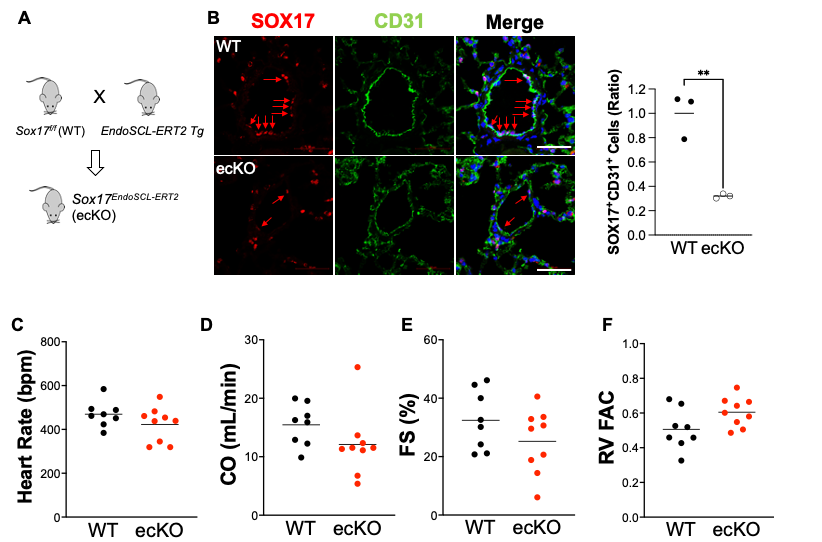

**Supplemental Figure 2. Generation of endothelial-specific knockdown of Sox17 in mice. (A)** A diagram showing the strategy of generating ecKO Sox17 mice. (B) Immunostaining demonstrated that SOX17 expression was reduced in the PVECs of ecKO Sox17 mice. Red arrows indicated SOX17 positive ECs. SOX17^+^ CD31^+^ cells in the pulmonary arteries from 10 fields were counted. The number of SOX17^+^ CD31^+^ cells from ecKO Sox17 mice was normalized by that from WT mice. (C-F) Echocardiography measurement showed that there were not significant change of heart rate, cardiac output (CO), fraction shorting (FS) and RV fraction area change (RV FAC) in ecKO Sox17 mice. Student t test. **, P< 0.01. Scale bar, 50$\mu$m.

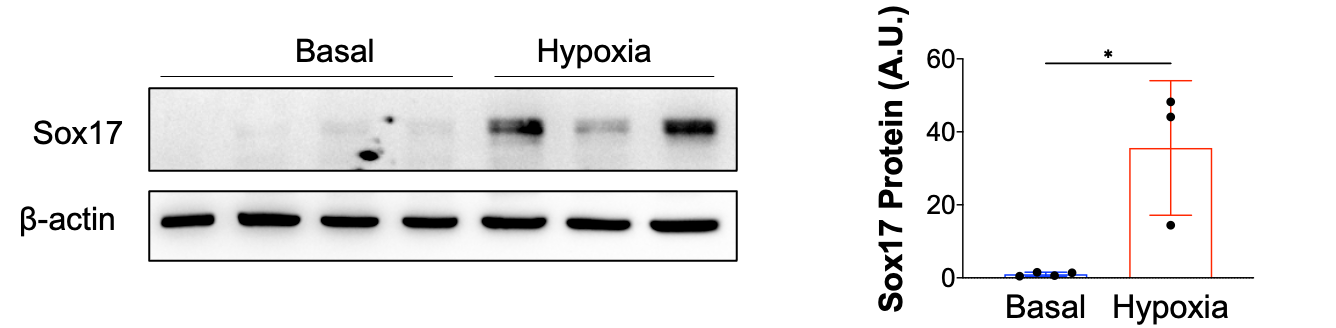

**Supplemental Figure 3. SOX17 expression was upregulated by hypoxia in mice.** Protein lysates were isolated from lung tissues from mice at 3 weeks post hypoxia treatment or similar age mice at the basal condition. Student t test. *, P< 0.05.

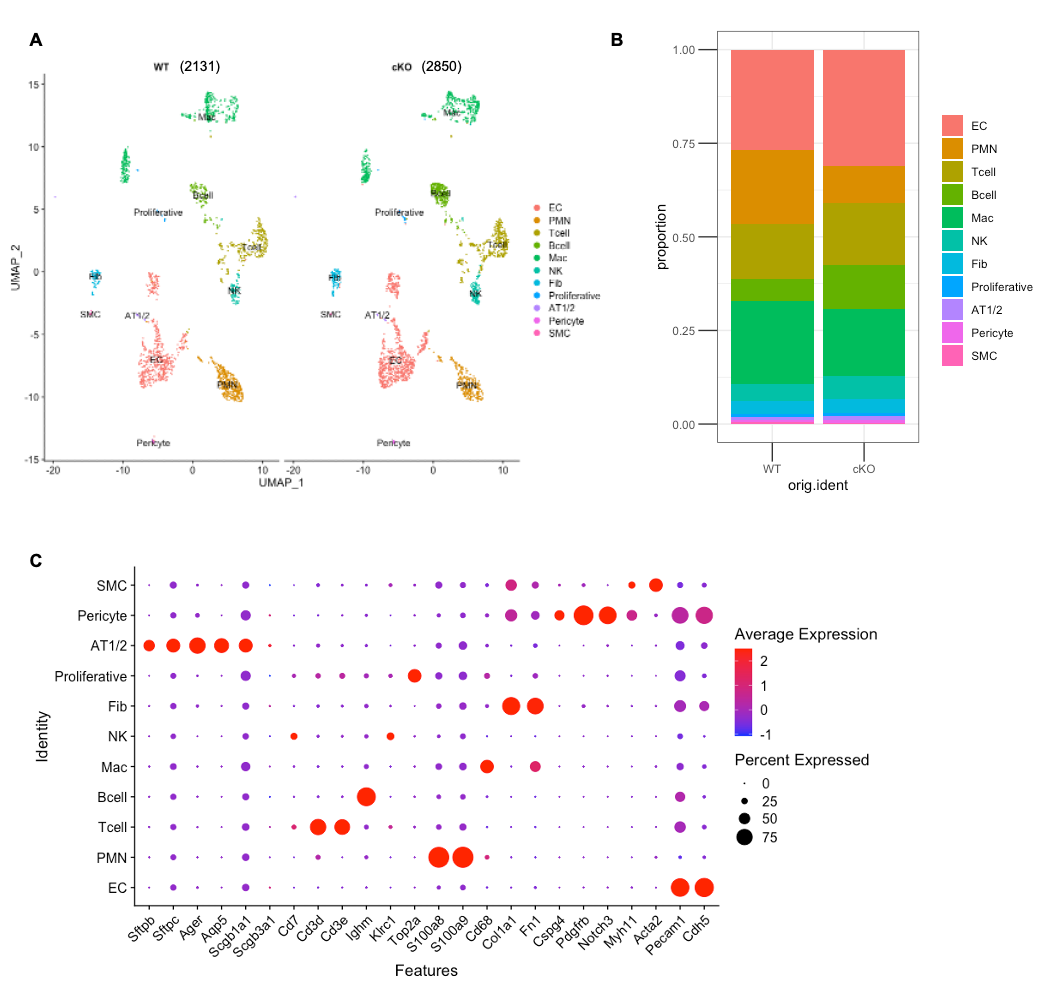

**Supplemental Figure 4. ScRNA-seq analysis of WT and cKO mice.** (A) A UMAP plot showing the clusters of lung cells. 2,131 and 2,850 cells were passed quality control from WT and cKO lungs. (B) Proportion of lung cells in A. There is an increase of EC, B cells population, and a reduction of PMN. (C) A Dot plot showing the lung ECs subpopulation based on their markers. PMN=neutrophil; Mac=macrophage/monocyte; NK=natural killer cell; Fib=fibroblast; AT1/2=alveolar type 1 and 2 epithelial cell; SMC=smooth muscle cells.

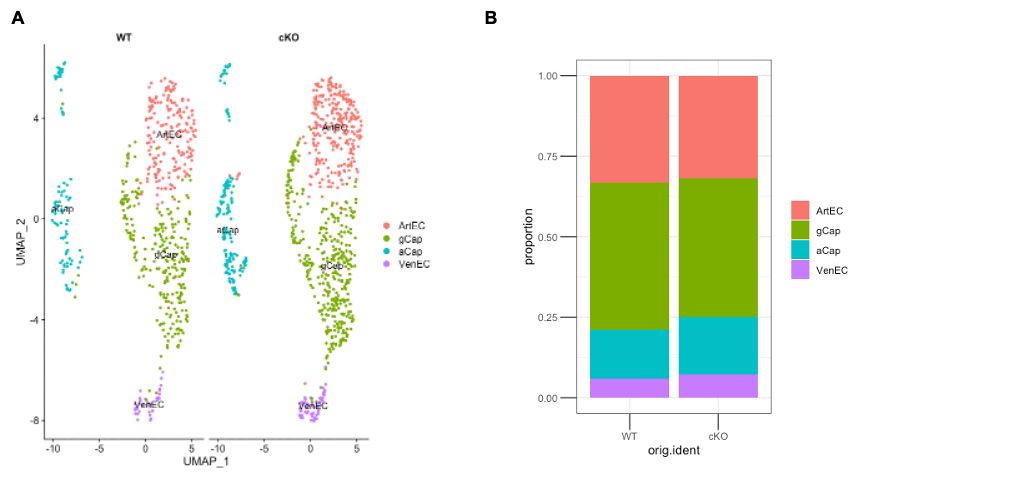

**Supplemental Figure 5. ScRNA-seq analysis of the ECs from WT and cKO mice.** (A) An UMAP plot showing the clusters of lung EC subpopulation. (B) Proportion of lung ECs in A. ArtEC=arterial ECs; gCap=general capillary; aCap=aerocyte; VenEC= venous EC.

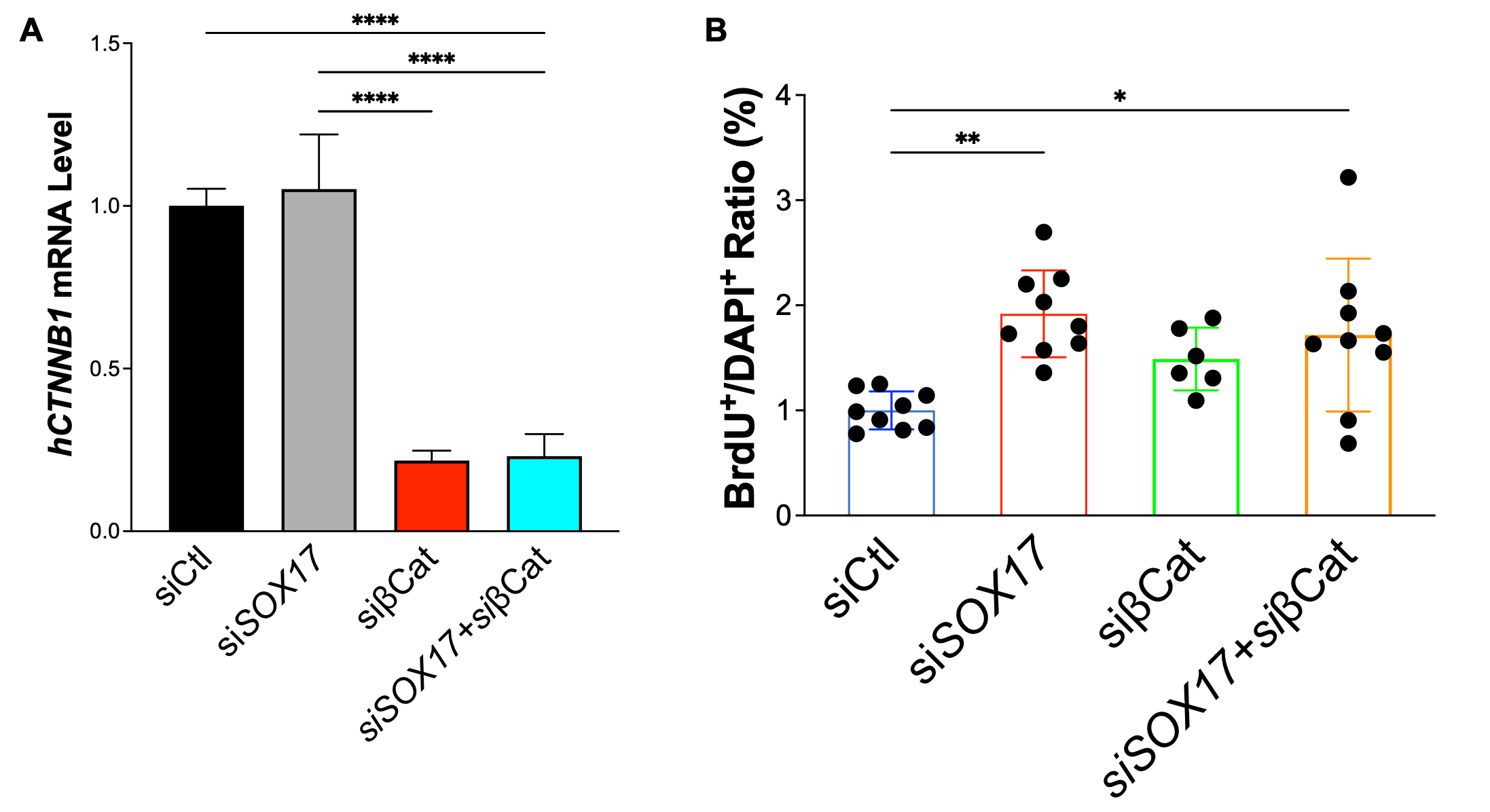

**Supplemental Figure 6.** β-catenin knockdown did not block the pro-proliferative effect induced by loss of SOX17. (A) QRT-PCR analysis showed that SOX17 and β-catenin were effectively knocked down. (B) BrdU incorporation assay demonstrated that β-catenin knockdown did not inhibit SOX17 deficiency induced EC prefoliation. One-way ANOVA with Tukey post hoc analysis. *, P< 0.05; **, P< 0.01, ****, P< 0.0001.

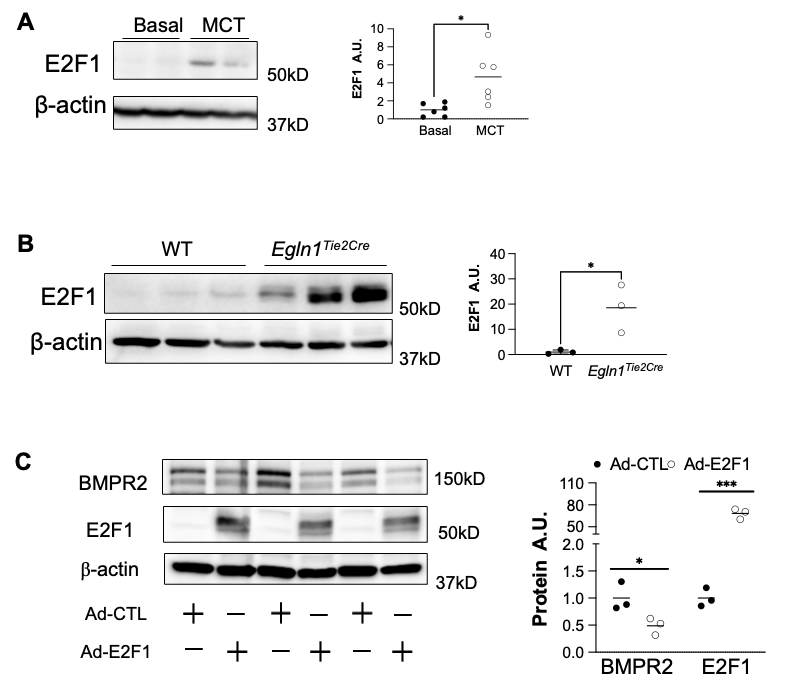

**Supplemental Figure 7. E2F1 is upregulated in the lung of MCT exposed rats and *Egln1^Tie2Cre^* mice. (A)** Protein lysates were isolated from lung tissues from rats at 4 weeks post MCT treatment or similar age rats at the basal condition. **(B)** Protein lysates were isolated from lung tissues from WT and *Egln1^Tie2Cre^* mice at the basal condition. **(C)** HPVECs were treated with adenovirus overexpressing E2F1 (10 M.O.I.) or control for 48 h. Student t test (A and B). *, P< 0.05.
